# SeSaMe: Metagenome Sequence Classification of Arbuscular Mycorrhizal Fungi Associated Microorganisms

**DOI:** 10.1101/2020.05.20.106559

**Authors:** Jee Eun Kang, Antonio Ciampi, Mohamed Hijri

## Abstract

Arbuscular mycorrhizal fungi (AMF) are plant root symbionts that play key roles in plant growth and soil fertility. They are obligate biotrophic fungi that form coenocytic multinucleated hyphae and spores. Numerous studies have shown that diverse microorganisms live on the surface and inside their mycelia, resulting in a metagenome when whole genome sequencing (WGS) data are obtained from sequencing AMF cultivated *in vivo.* The metagenome contains not only the AMF sequences, but also those from associated microorganisms. In this article, we introduce a novel bioinformatics program, SeSaMe, designed for taxonomic classification of short sequences obtained by next-generation DNA sequencing. A genus-specific usage bias database was created based on amino acid usage and codon usage of three consecutive codon DNA 9-mers encoding for an amino acid trimer in a protein secondary structure. The program distinguishes between coding sequence (CDS) and non-CDS, and classifies a query sequence into a genus group out of 54 genera used as reference. The average correct prediction percentages of the CDS and the non-CDS test sets at the genus level were 71% and 50% for bacteria, 65% and 73% for fungi (excluding AMF), and 49% and 72% for AMF (*Rhizophagus irregularis*), respectively. The program provides a means for estimating not only taxonomic diversity and abundance but also the gene reservoir of the reference taxonomic groups associated with AMF. Therefore, the program enables users to study the symbiotic roles of associated microorganisms. SeSaMe can be applicable to other microorganisms as well as soil metagenomes. It is freely available at www.journal.com and www.fungalsesame.org.

## Introduction

Arbuscular mycorrhizal fungi (AMF) are plant root inhabiting fungi, of the subphylum Glomeromycotina, which form symbioses with more than 80% of vascular plants worldwide [1]. They supply plants with essential nutrients particularly phosphorus and nitrogen, protect them against soil borne pathogens, and alleviate their abiotic stresses [1–3]. Therefore, AMF based inoculants have been applied in agriculture as a biofertilizer and in phytoremediation for cleaning up contaminated soil [2,4–7]. Despite the ecological, agricultural, and environmental importance of AMF, their genetics is poorly understood due to their complex genome organization. They form coenocytic hyphae, reproduce through multinucleated asexual spores, and are strict symbionts [8]. Furthermore, it is suggested that AMF are heterokaryons, although this is under debate [9]. In addition, numerous studies reported that bacteria and fungi inhabit the surface and the interior of mycelia and spores [10–14]. In 2012 and 2013, Tisserant et al. published the transcriptome and the genome of the AMF *Rhizophagus irregularis* cultivated *in vitro* [15,16]. However, only a few AMF taxa are able to grow in axenic *in vitro* systems with transformed roots as a host. Thus, whole genome sequencing (WGS) data from AMF spore DNA originating from *in vivo* cultures (conventional cultivation method in a pot culture with a host plant), contain a substantial number of non-AMF DNA sequences, but do provide important information on the microbial communities associated with AMF. In contrast, WGS data from *in vitro* petri-dishes contain fewer non-AMF sequences, because antibiotics are used to initiate axenic cultures [17].

Taxonomic classification of WGS obtained from AMF cultivated *in vivo* using current bioinformatics approaches is challenging because these data represent a complex metagenome containing sequences of prokaryotic and eukaryotic microorganisms. Two major approaches for taxonomic classification of random whole metagenome sequencing data (*e.g.*, whole metagenome shotgun sequencing data) include composition-based methods and similarity-based search methods [18,19]. The latter ones include BLAST and its sister programs that are adequate for inferring functions of a query sequence [19,20]. Nevertheless, they have limitations in taxonomic classification, because they calculate scores based on a 20 by 20 matrix containing the overall rates of the 20 amino acid substitutions created from the most conserved regions of proteins. The same matrix is applied to all types of query sequences, irrespective of functions, structures, and taxonomic group. However, due to a lack of bioinformatics tools for analyzing random whole metagenome data, similarity-based search methods have been commonly used for taxonomic classification. In addition to similarity-based methods, taxonomic classification pipelines, for analyzing targeted metagenome sequencing data (*e.g.*, 16S rRNA gene-based metagenome sequencing data), have been widely used for analyzing random whole metagenome sequencing data in combination with homology search program. Numerous repository databases and pipelines have been developed based on the 16S rRNA gene. However, recent studies have reported horizontal gene transfer of 16S rRNA genes in prokaryotic organisms and multiple heterogeneous rRNA genes within a single prokaryotic cell [21]. Therefore, they may cause misrepresentation of data if they are not properly dealt with, which may result in erroneous taxonomic classification.

Composition-based methods utilize unique sequence properties such as codon usage bias, compositional patterns in nucleotide sequences (k-mers), and GC content that have been widely used for studying microbial genome evolution in areas of bioinformatics [18,22–26]. K-mers are subsequences of length k in a DNA sequence (*e.g.*, tetramer or 4-mer: ATGT). Composition-based methods using k-mers have been employed in bioinformatics programs for taxonomic classification of random whole metagenome data [27]. They have a number of advantages over similarity-based search methods. It is estimated that more than 99% of existing microorganisms cannot be cultured in laboratory conditions [28] and microbial sequences available in bioinformatics databases represent only a tiny fraction of the diversity of existing microorganisms. Therefore, composition-based methods, that do not require sequence alignments but make predictions based on a microorganism’s unique sequence signatures, supposedly excel in taxonomical classification of novel sequences. However, existing bioinformatics programs based on composition-based methods are designed for prokaryotic organisms and their utilization in fungi is inefficient.

In this article, we introduce a novel bioinformatics program for random whole metagenome sequence classification, SeSaMe (Spore associated Symbiotic Microbes). It provides a means for estimating taxonomic diversity and abundance, as well as, the reservoir of genes of reference taxonomic groups in AMF metagenome. It therefore enables users to study symbiotic roles of taxonomic groups associated with AMF. In order to filter complex evolutionary signals and obtain comparable evolutionary footprints, we calculated codon usage bias based on the amino acid usage and the codon usage of three codon DNA 9-mer that encodes for three consecutive amino acids located in protein secondary structure. We joined three consecutive codons into one unit, and calculated the unit’s relative frequency among synonymous three codon DNA 9-mers, which will be hereafter referred to as three codon usage. Three codon usage has higher resolution than mono codon usage in assessing the differences among taxonomic groups because evolutionary forces acting on a codon and its encoded amino acid vary widely across protein secondary structures as well as across taxonomic groups. For example, the evolutionary forces acting on the codon AAA, encoding the amino acid Lysine (K) in TGG***AAA***GTG (WKV), will have been different from the evolutionary forces acting on the codon AAA in GAC***AAA***GAA (DKE). We found that three codon usage of a three codon DNA 9-mer belonging to protein secondary structure is a taxonomically unique sequence property. SeSaMe calculates a score based on six sets of three codon DNA 9-mers from all reading frames (**Figure 1**), and distinguishes between coding sequence (CDS) and non-CDS. It has an advantage over existing composition-based methods that do not identify nucleotide subsequences with structural roles, or do not consider the biological importance of codon and reading frame. SeSaMe is freely available at www.journal.com and www.fungalsesame.org and www.fungalsesame.org.

**Figure 1.**
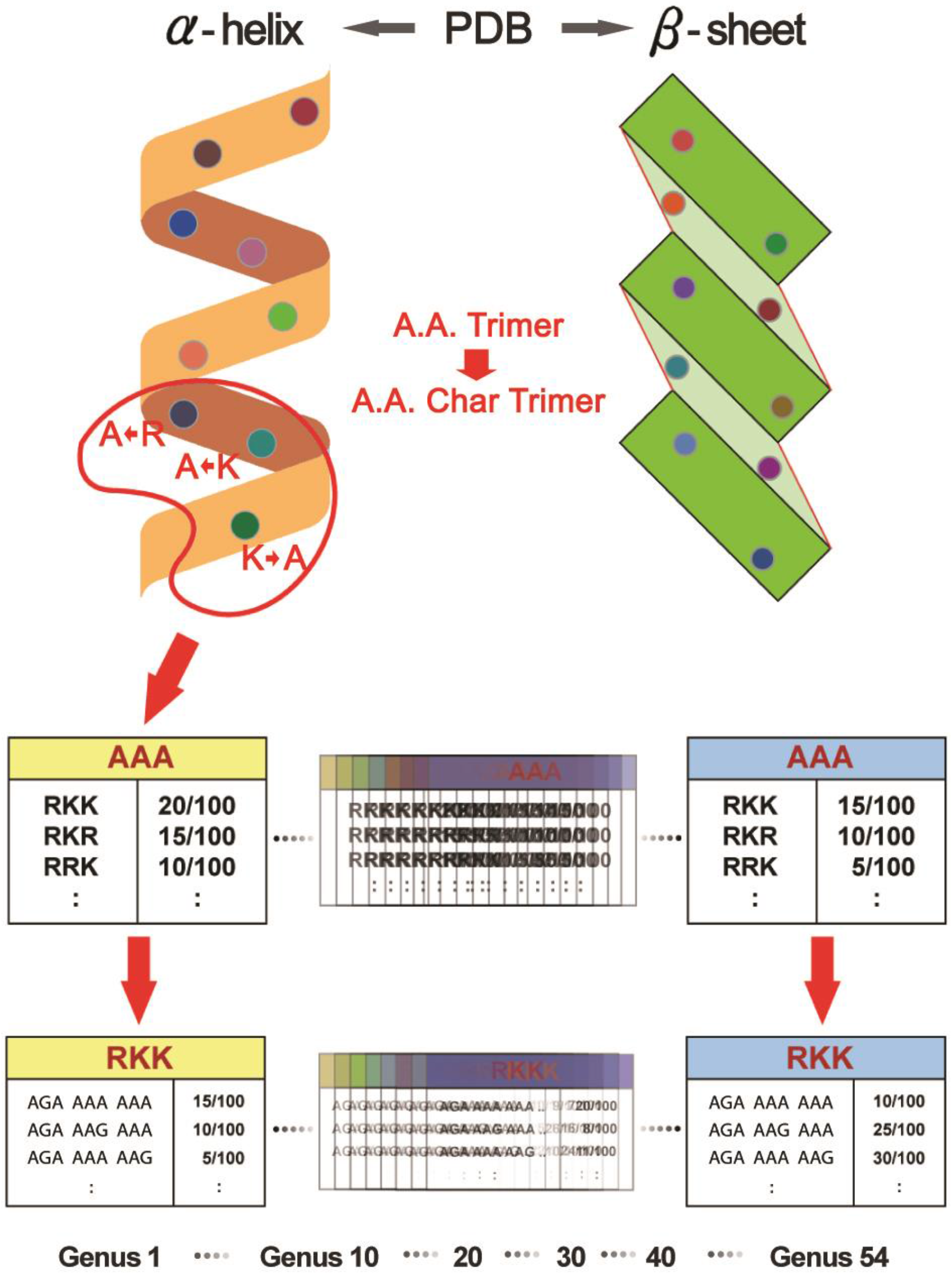
Unique advantage of SeSaMe over existing programs. Existing programs calculate a score based on the frequencies of k-mers identified in a query sequence irrespective of properties of the k-mers, or its reading frame. In contrast, SeSaMe identifies k-mers that encode for the amino acids of protein secondary structures in each reading frame. In the figure, matching Three Codon DNA 9-mers of the Trimer Ref. DB are marked with a rectangle, where the rectangle’s color indicates its reading frame. The program calculates scores based on the three codon usages and the A.A. Trimer usages of the matching Three Codon DNA 9-mers in each reading frame. It classifies a query sequence into a taxonomic group based on the six scores computed from all reading frames. All sequences in this figure are randomly generated for illustration purposes only.

## Methods

### Bacterial and fungal sequence databases

We selected bacterial genera that were dominant in soil based on a literature review [10, 29,33–36]. While NCBI offered a broad selection of more than 2,300 completely sequenced bacterial genomes, we did not have many choices for the majority of fungal phyla. Most of the completely sequenced fungal genomes in NCBI or JGI were Dikarya, while we needed diverse fungal genomes covering Mucoromycotina, AMF, Blastocladiomycota, Neocallimastigomycota, Microsporidia, and Chytridiomycota. We assigned the completely sequenced genomes of 444 bacteria and of 11 fungi, including *R. irregularis*, to 45 bacterial and 9 fungal genera respectively, and created CDS and non-CDS databases per genus based on CDS lists provided by NCBI, JGI, and Tisserant et al. [16]. The number of genomes per genus varied from 1 to 81, depending on their availability in public databases. The total number of the bacterial, and the fungal, genes and introns, per genus, are shown in Tables S1 and S2. Sequences with an ambiguous nucleotide or with a length shorter than nine—the minimum length of nucleotides required for three codon DNA 9-mers—were excluded. *Cryptococcus* and Agaricomycetes (*Phanerochaete*, *Scleroderma, Sebacina*) belong to the same subdivision, Agaricomycotina, and were grouped together in order to simplify the analysis.

### Database design

In selecting a parameter k of k-mer, we chose three codon DNA 9-mer as the length of amino acid and of nucleotide, considering the approximate number of amino acids required to form a turn in helix and a beta-strand. The program consists of two main components: databases and scoring methods. The major distinguishing feature is the trimer reference sequence database (Trimer Ref. DB). 126,093 Protein Data Bank (PDB) entry files were processed with in-house developed parsing programs in order to extract 7674 amino acid trimers, subunits of protein secondary structures, which were assigned to the sequence variable-A.A. Trimer [33]. 224,383 three codon DNA 9-mers, encoding 7674 A.A. Trimers, were assigned to the sequence variable-Three Codon DNA 9-mer. In Trimer Ref. DB, the sequence variables-A.A. Char Trimer, A.A. Trimer, and Three Codon DNA 9-mer form a three level hierarchy where A.A. Char Trimer is the highest level (**Figure 2**). To create amino acid characteristic (A.A. Char), first, we assigned amino acids with similar properties into one group according to polarity and charge of their side chain, and secondly subdivided each group according to their volume (**Table 1**). Cysteine, Glycine, Histidine, Methionine, and Proline have special properties; Cysteine forms disulfide bonds, Glycine is the simplest amino acid, Histidine can be a proton shuttle, Methionine is often the first amino acid, Proline is animino acid. Therefore, each of them was assigned as a sole member of A.A. Char group. Generally, multiple A.A. Trimers with similar properties belong to one A.A. Char Trimer. An A.A. Char Trimer and an A.A. Trimer have A.A. Trimer table and Three Codon DNA 9-mer table containing multiple members, respectively (Figure 2).

**Table 1.** Conversion table from A.A. to A.A. Char.

| A.A. Char | A.A. | Properties | A.A. Char | A.A. | Properties |
| --- | --- | --- | --- | --- | --- |
| A | K,R | Positively charged | G | G | Special |
| B | H | Special | H | P | Special |
| C | D,E | Negatively charged | I | M | Special |
| D | S,T | Polar uncharged smaller size | J | A,I,L,V | Hydrophobic smaller size |
| E | N,Q | Polar uncharged larger size | K | F,W,Y | Hydrophobic larger size |
| F | C | Special | L | * | Stop codon |

*Note:* Amino acids were grouped according to their side chain's pKa values and charges at physiological pH (7.4) and sizes (www.wikipedia.org/amino_acid)

**Figure 2.**
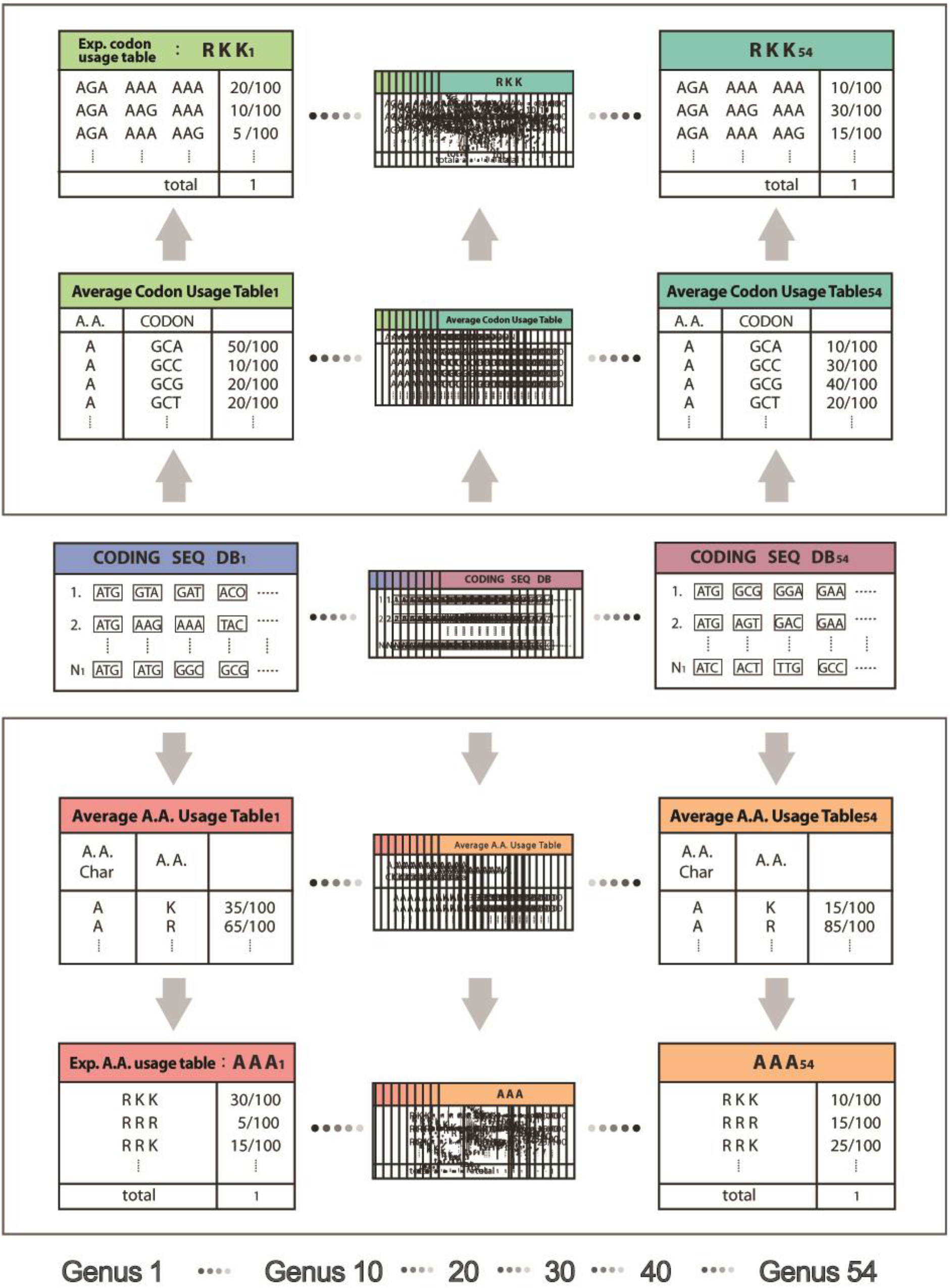
Database design. In this figure, A.A. Trimer Usage Table consists of the A.A. Trimer usages of the multiple members-RKK, RKR, and RRK belonging to the same A.A. Char Trimer-AAA. Three Codon Usage Table consists of the three codon usages of the synonymous Three Codon DNA 9-mers encoding the A.A. Trimer, RKK (*e.g.*, AGA AAA AAA). The trimer usage bias of AGA AAA AAA is the multiplication of the A.A. Trimer usage of RKK and the three codon usage of AGA AAA AAA. All sequences and usage information in this figure are not real, but randomly chosen for illustration purposes only.

Genus-specific usage bias database (Genus Specific DB) contains the main numerical variable, trimer usage bias. Trimer usage bias represents a three codon usage bias of Three Codon DNA 9-mer, and is calculated by multiplying the A.A. Trimer usage of A.A. Trimer by the three codon usage of Three Codon DNA 9-mer in Trimer Ref. DB (Figure 2). There are 54 CDS Genus Specific DBs and the same number of non-CDS Genus Specific DBs in the program. Each CDS Genus Specific DB contains 1296 A.A. Trimer Usage Tables and 7674 Three Codon Usage Tables created based on the CDS database. Each non-CDS Genus Specific DB contains the same number of tables created based on the non-CDS database with the same sequence compositions as those in the CDS Genus Specific DB. We decided to accept inaccuracy in calculating the information frequency in the case of non-CDS in exchange for cost effective CDS and non-CDS classification. Because SeSaMe only needs to compare frequency information of 54 genera calculated based on the same standard genetic code table for the same Three Codon DNA 9-mers of a query sequence, inaccuracy in non-CDS is assumed to be insignificant.

### Scoring methods

We developed two scoring methods, and each equipped with a *P* value scoring method. The trimer usage probability scoring method classifies a query sequence into one out of 54 genus references, while the rank probability scoring method classifies a query sequence into one out of 13 taxon groups: Clostridia, Bacilli, Oscillatoriophycideae, Nostocales, Acidobacteriales, Betaproteobacteria, Deltaproteobacteria, Gammaproteobacteria, Alphaproteobacteria, Actinobacteria, AMF (*R. irregularis*), Agaricomycotina, and Pezizomycotina. To avoid repetition, these taxonomic groups will be hereafter referred to as 13 taxon groups, and represented in the same order. We provide users with two different programs, one with the trimer usage probability scoring method and the other with the rank probability scoring method.

#### Trimer usage probability scoring method

This method converts three codon DNA 9-mers in a query sequence into A.A. Char Trimers and identifies those with structural roles by searching them against Trimer Ref. DB. For each matching A.A. Char Trimer, the method first searches the matching A.A. Trimer, and second, the matching Three Codon DNA 9-mer in Trimer Ref. DB (**Figure 3**). It retrieves trimer usage biases of the matching Three Codon DNA 9-mers from CDS Genus Specific DB per genus. It repeats the process in each of 6 reading frames (3 forward reading and 3 reverse reading frames) of a query sequence. It repeats the same process with non-CDS Genus Specific DBs, calculating a trimer usage probability score per genus. It then compares the highest scores from CDS and non-CDS Genus Specific DBs, and selects a genus with the highest score (Figure 3). Users are provided with an option to include genera whose scores have little difference from the highest score calculated.

**Figure 3.**
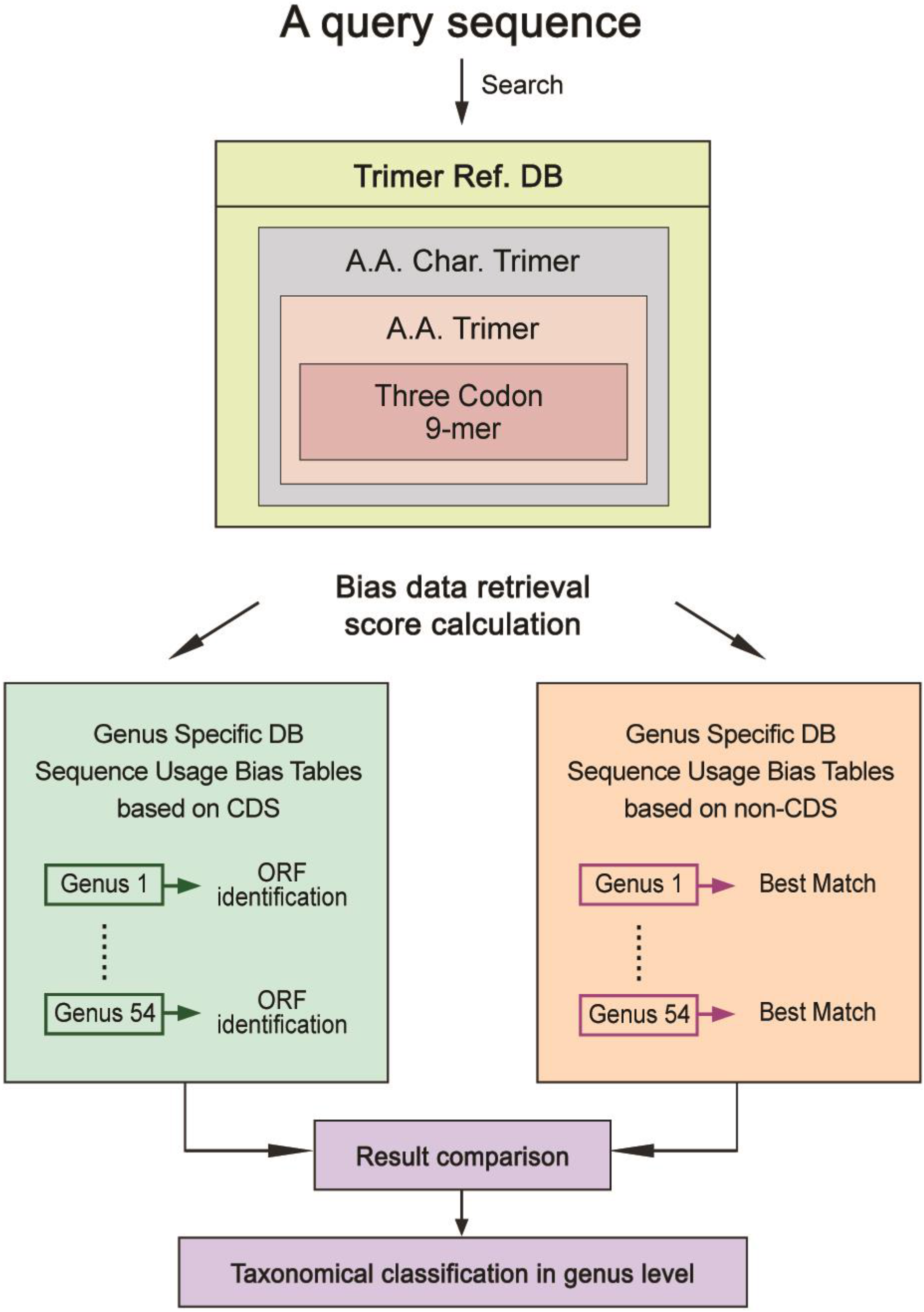
Flow chart of the program.

#### Rank probability scoring method

This method measures a standardized three codon usage relative to an expected three codon usage as computed from three individual mono codon usages. The Average A.A. Usage Table (20 amino acids and stop codons for 12 A.A. Char monomers) and the Average Codon Usage Table (64 codons for 20 amino acid monomers and stop codons) were created based on CDS database per genus. 1296 Expected A.A. Trimer Usage Tables with the same sequence compositions as the A.A. Trimer Usage Tables were created based on the Average Amino Acid Usage Table. 7674 Expected Three Codon Usage Tables with the same sequence compositions as the Three Codon Usage Tables were created based on the Average Codon Usage Table (**Figure 4**).

**Figure 4.**
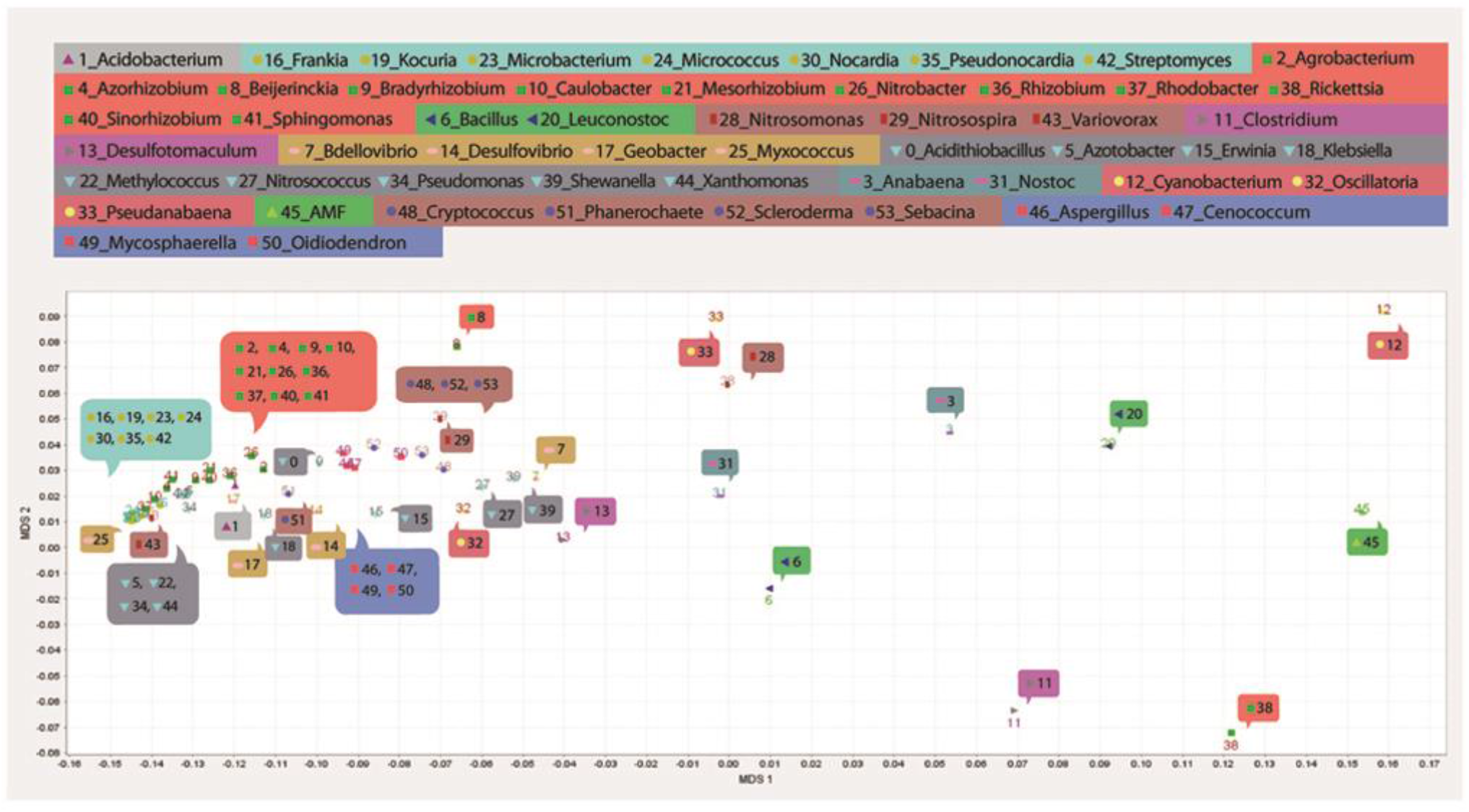
Creation of expected usage tables for the rank probability scoring method. Average A.A. Usage Table and Average Codon Usage Table were calculated from the CDS database per genus. Expected A.A. Trimer Usage Tables and Expected Three Codon Usage Tables were created based on the Average A.A. Usage Table and the Average Codon Usage Table, respectively. All sequences and expected usage information in this figure are not real, but randomly chosen for illustration purposes only.

A standardized three codon usage was calculated by dividing a three codon usage in a Three Codon Usage Table by an expected three codon usage in an Expected Three Codon Usage Table. Based on trimer usage biases and standardized three codon usages, we calculated a group mean for each taxon group and Kruskal Wallis (KW) test’s h-score on ranks of 13 taxon groups, from which we developed a rank probability score per Three Codon DNA 9-mer. The new genus specific score database contains the same number of the rank probability scores as Genus Specific DB, 224,383 scores per genus. For per reading frame of a query sequence, the program retrieves scores for all of the matching Three Codon DNA 9-mers from the new database and multiplies the scores to produce a rank probability score per genus. It repeats the process for each of 6 reading frames and classifies a query sequence into one of 13 taxon groups. This method is applicable only to CDS.

#### P value scoring method

We applied the concept of the sum of rolled numbers from a pair of dice to develop the *P* value scoring method (http://www.lucamoroni.it/the-dice-roll-sum-problem/). We drew analogies between the number of faces of a dice and 54 genera and between the number of dices we roll and the number of matching Three Codon DNA 9-mers identified in a reading frame of a query sequence. There were 54 possible ranks computed based on trimer usage biases per matching Three Codon DNA 9-mer. *P* value scores were calculated based on a sum of ranks of matching Three Codon DNA 9-mers. Computational costs of *P* values for all possible outcomes, sums of ranks, were too high, however, so to reduce the computational costs we approximated *P* values. We obtained sample data per number of matching Three Codon DNA 9-mers based on equation 1.

Equation. 1:

where *p* is the sum of ranks, *n* is the number of dices per roll, *s* is the number of faces of the dice, 54, and maximum of *k* is ](*p*-*n*)/*s*[ where ]*x* [is the floor function (*e.g.*, ]7.9[ = 7). We created a table of *P* value scores per number of matching Three Codon DNA 9-mers. If a rank sum was less than one with the highest *P* value score, the approximate mean of all of the rank sums in each table, we multiplied the *P* value score with −1, indicating statistically non-significant outcome. In the test sets, the number of matching Three Codon DNA 9-mers varied widely, with a minimum of 30 and a maximum of 97. We have 624 tables in the *P* value score database covering 2–625 matching Three Codon DNA 9-mers. *P* value scores are generated per genus in both the trimer usage probability, and the rank probability, scoring methods to provide users with the statistical significance of predicted outcomes.

### Implementation and program availability

SeSaMe has been implemented using the Java programming language (www.java.net, www.oracle.com (Java 8)). We have provided two sets of the programs; one requires Apache commons math3 (3.3) and IO (2.4) libraries (www.apache.org), while the other does not. The programs consist of executable Java JAR files and Java class files for Linux/Unix operating systems. SeSaMe has been tested and confirmed to work on Linux system−CentOS Linux 7 (www.centos.org) and is currently being used at the Biodiversity Center, Institut de Rechercheen Biologie Végétale, Département de Sciences Biologiques, Université de Montréal. The trimer usage probability scoring method offered to the public produces output of smaller size, but is sufficient for the purpose of taxonomic classification and is freely available at www.journal.com and www.fungalsesame.org. There are no restrictions to use the programs by academic, or non-academic, organizations as long as they comply with the terms and conditions of the license agreements.

### Input, output, and options

SeSaMe utilizes a command-line interface. Input files should contain DNA sequence(s) in fasta format. The Java JAR files produce detailed output files with sequence information (seq_id, matching A.A. Char Trimers, A.A. Trimers, and Three Codon DNA 9-mers) and genus information (rank, scores, and *P* value score). The output details the information per reading frame per sequence. After processing the output file with Java class files, users are able to obtain a summary file containing one predicted outcome per query sequence. Java JAR files require users to give a mandatory command line argument-input file path. Java JAR files with the trimer probability scoring method may produce multiple genera as an answer if their scores have little differences. A user is given the option with 6 choices to select a cut-off value for the difference: 0.01, 0.05, 0.1, 0.15, 0.2, or 0.3. Users can give the option to the Java class file called compare_result_coding_non_coding.class. The default cut-off value is 0.05. The lower the cut-off value is, the fewer genera will be included in an answer.

### Program evaluation

We assessed the accuracy of the classification program by conducting classification experiments. We created metagenome test sets, ran the programs with them, and calculated the correct prediction percentages. We showed the relationship between the correct prediction proportion and the *P* value score in order to provide users with useful examples in assessing the statistical significance of predicted outcomes.

#### Metagenome test sets

We randomly chose 100 sequences from each of the CDS and non-CDS databases for each genus. We randomly selected a starting base pair position in each of the chosen sequences. From the starting position, we randomly selected an ending base pair position so that a sequence length is within the range of 150–300 bp. Both of the CDS and the non-CDS test sets consisted of 4500 bacterial and 900 fungal sequences (including AMF).

#### Correct prediction percentages from the trimer usage probability scoring method

The means of the correct prediction percentages of the CDS and the non-CDS test sets at the genus level were 71% and 50% for the bacterial group, 65% and 73% for the fungal group (excluding AMF), and 49% and 72% for AMF, respectively. AMF showed the lowest prediction percentage among the CDS genus test sets possibly due to a large number of heterogeneous nuclei and horizontal gene transfers from a variety of endobacteria during their evolution [8,10–14]. The means of correct prediction percentages at the genus level and at higher taxonomic ranks of the 13 taxon groups are shown in **Table 2**.

**Table 2.**
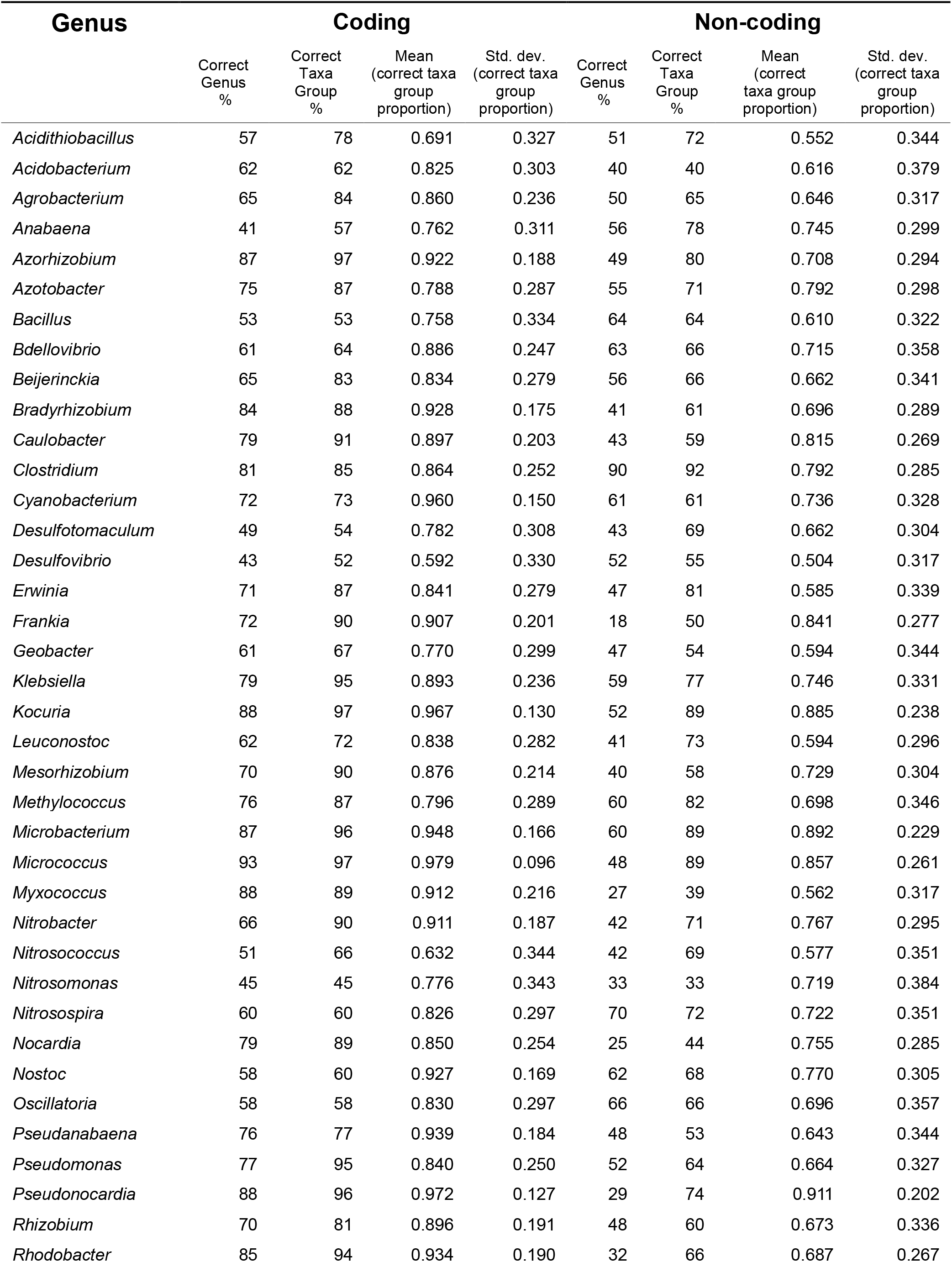
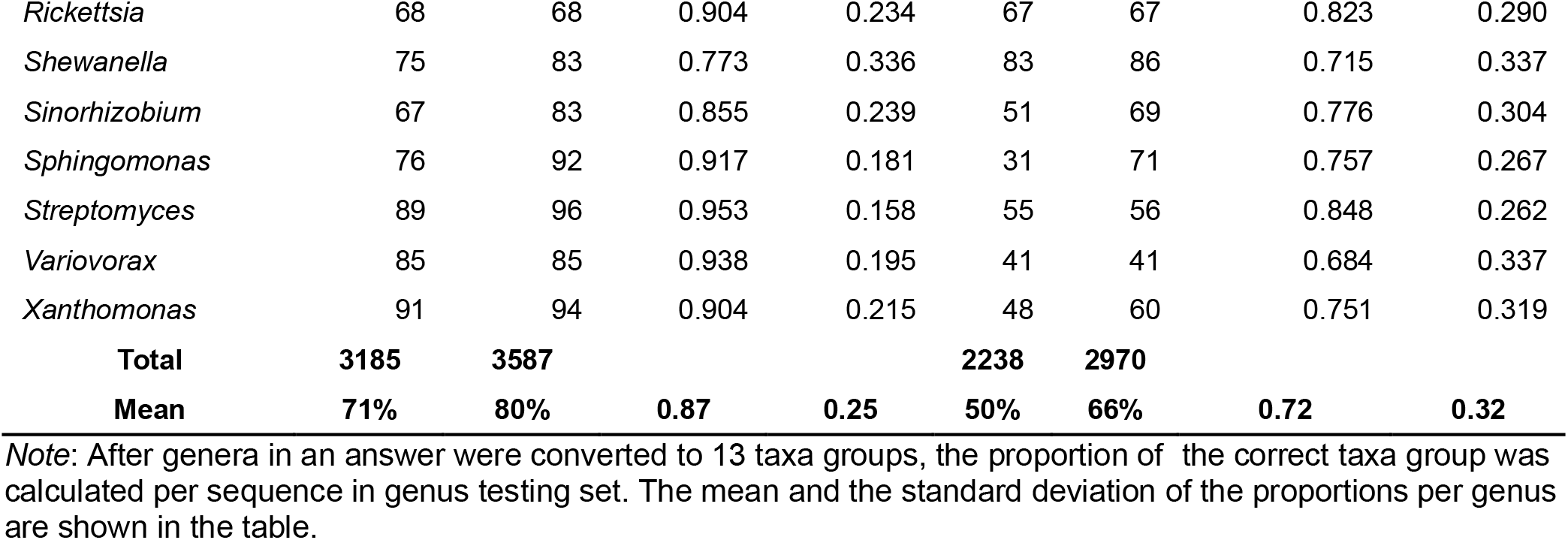
Bacterial correct prediction percentages in 54 genus and 13 taxa group levels.

SeSaMe produced more than one genus as an answer per query sequence when multiple genera had little differences in their scores. We converted each predicted genus into one of the 13 taxon groups and calculated a proportion of the correct taxon group in answer per query sequence. We calculated the mean and the standard deviation of the proportions in each genus test set; 1 represented that answers contained correct taxon groups only, while 0 represented that answers contained incorrect taxon groups only (Tables S3 and S4). The mean was 0.9 for the bacterial CDS test set, which indicated that in average 90% of the taxon groups in an answer were correct.

SeSaMe produced only one genus as an answer in 60% and 46% of correctly predicted sequences from the bacterial CDS and non-CDS test sets, respectively (Figure S1, Table S5). A correct taxon group occurred in the first rank in 90% and 76% of correctly predicted sequences in the bacterial CDS and non-CDS test sets, respectively (Figure S2, Table S6). Only 1% – 5% of the sequences in the bacterial and the fungal test sets had AMF in an answer (Table S7). Although the trimer usage probability scoring method provides us not with the individual trimer usage biases but with the result of multiplying all of the trimer usage biases identified in a query sequence, we can often derive general ideas about the query sequence from its answer. Does it contain only one genus in the answer? Or what other genera does it contain in answer? (Figures S3 and S4) For example, an AMF test sequence that contains *Clostridium* and AMF in the answer may imply that the query sequence may have been acquired by horizontal gene transfer from a bacterium ancestor to an AMF ancestor during evolution.

#### Correct prediction percentages from the rank probability scoring method

The means of the correct prediction percentages of the CDS test set were 82% for the bacterial group, 72% for the fungal group (excluding AMF), and 42% for AMF. The means and the standard deviations of the correct prediction percentages of the CDS test set were 64% ± 4.2%, 71% ± 6.4%, 84% ± 2.5%, 70% ± 2.8%, 73% ± 0%, 83% ± 8%, 74% ± 10%, 81% ± 7.8%, 88% ± 9.2%, 85% ± 5.9%, 42% ± 0%, 65% ± 6.4%, and 79% ± 6.7% for the 13 taxon groups, respectively. Compared to the trimer usage probability scoring method, the rank probability scoring method produced the higher mean and the smaller standard deviation, 82% ± 9.4% for the bacterial group. In general, the rank probability scoring method showed improvement in performance. Although the means for Clostridia and Gammaproteobacteria were lower, their standard deviations were much smaller in the rank probability scoring method than the trimer usage probability scoring method: 4.2% *vs* 22% and 7.8% *vs* 9.4%, respectively. The trimer usage probability scoring method showed better performance in Actinobacteria that had low within-group variation of trimer usage bias. In contrast, the rank probability scoring method showed better performance in genera that had relatively flat peakness in a frequency distribution curve of synonymous Three Codon DNA 9-mers, in addition to genera that had relatively large within-group variation of trimer usage bias.

#### Relationship between correct prediction proportion and P value score

The means of the correct prediction proportions per number of matching Three Codon DNA 9-mers calculated based on the result of the trimer usage probability scoring method is shown in Figure S5A and Table S8. The means of correct prediction proportions per base 10 logarithm of an approximated inverse of a rank sum based *P* value score (log10 (inverse of *P* value score)) calculated based on result of the trimer usage probability scoring method and of the rank probability scoring method are shown in Figure S5B, Table S9, Figure S5C, and Table S10, respectively. We divided the results of each genus test set into quartiles and calculated the range of (log10 (inverse of *P* value score)), the mean and the standard deviation of the correct prediction proportions in each quartile. They are shown in Tables S11 and S12 for the trimer usage probability scoring method and the rank probability scoring method, respectively. The first ranked genus with the highest probability score that was selected as an answer of a test sequence always had positive *P* value score. In general, as (log10 (inverse of *P* value score)) became higher—*i.e.*, as positive *P* value score became lower—the correct prediction proportion increased in all test sets. The frequencies of fungal sequences that had a correct taxon group in the 1st, 2nd, 3rd, 4th, or 5th rank, in an answer were comparable due to similarity of Dikarya (Figure S2, Table S6). Because the data for Figure S5B were generated based only on the first rank, the fungi showed relatively weak correlation between correct prediction proportion and (log10 (inverse of *P* value score)). The AMF database contains only one species, *R. irregularis*, therefore, results from both methods showed little difference.

#### Classification of an example sequence

Here we demonstrate the analysis of a query sequence selected from the AMF CDS test set. The example sequence was 156 bp (AAATCCCAATGTCAGAATAAAGAAACTACCAGATGATCATCCTGTTTATCCTGGGTATGGATTATTTGCTAACAAAGATCTTAAAAAATTTAATCTAGTCGTTTGTTATACTGGCAAAGTTACAAAAAGAGAAATTGGGGGTGAAGAAGGAAGTGA). The sequence had the highest trimer usage probability score in the second reading frame translation, which was then assumed as the open reading frame. SeSaMe identified 49 matching Three Codon DNA 9-mers in the second reading frame that were matched to Trimer Ref DB. The program correctly classified the example sequence into CDS of AMF. Firmicutes, Cyanobacteria, *Rickettsia*, and AMF had higher trimer usage biases than Proteobacteria, Actinobacteria, and Dikarya in a majority of Three Codon DNA 9-mers. **Figure 5** shows trimer usage biases of the 11th Three Codon DNA 9-mer-GATGATCAT in 54 genera.

**Figure 5.**
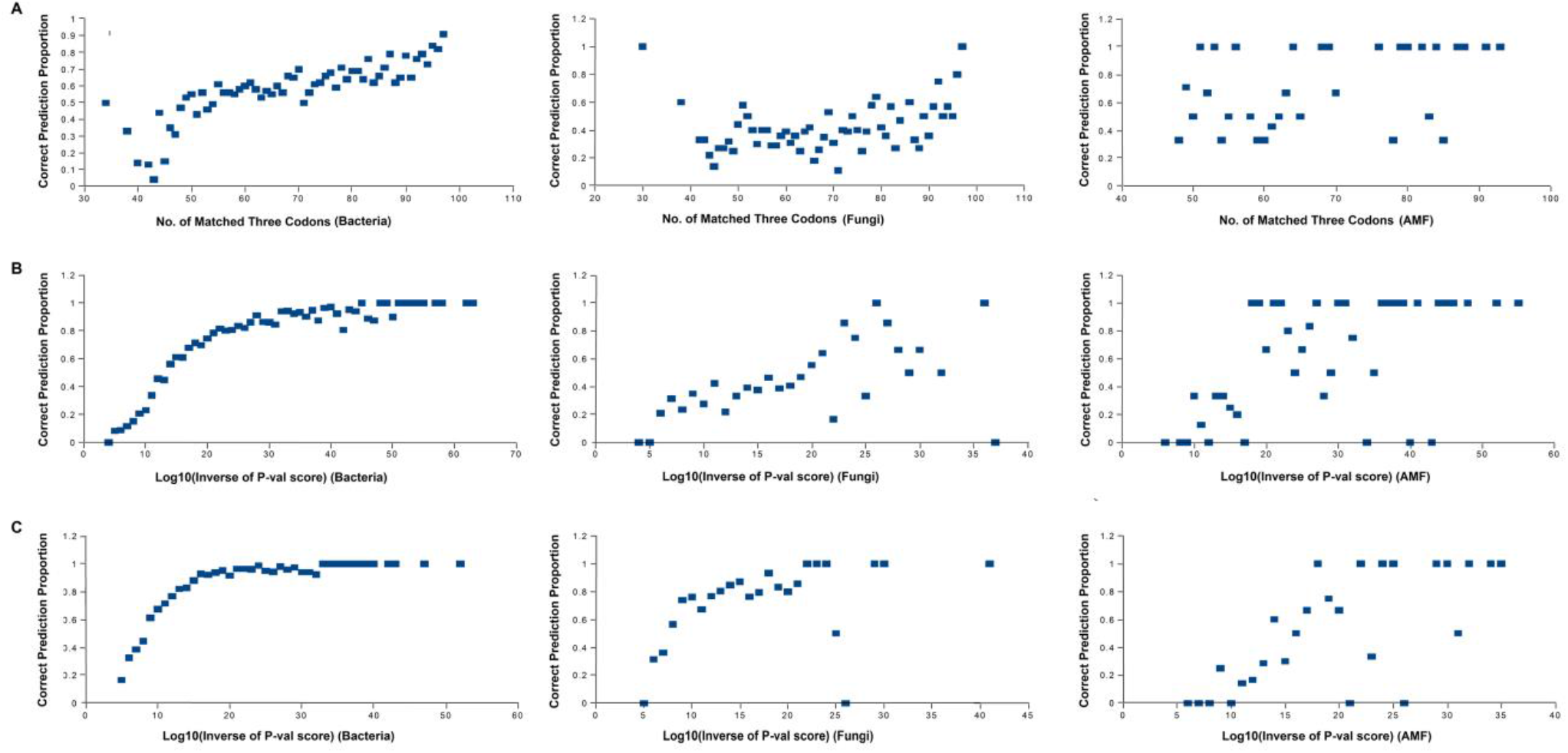
Trimer usage biases of 11_th_ Three Codon DNA 9-mer, GATGATCAT, in 54 genera. Genera belonging to the same taxonomic group are indicated by the same background color.

GATGATCAT belongs to A.A. Char trimer-CCB and to A.A. Trimer-DDH. The multidimensional scaling (MDS) method was applied to a matrix containing trimer usage biases; it had 54 genera in rows and matching Three Codon DNA 9-mers identified in the open reading frame in columns (http://www.inf.uni-konstanz.de/algo/software/mdsj/) [37]. It visualized proximity relationships among 54 genera in XY axis graph (www.jfree.org). It showed that Actinobacteria, Alphaproteobacteria, and Dikarya were compactly clustered, while Betaproteobacteria were spread out in the left side of the graph (Figure S6). Nostocales, Oscillatoriophycideae, Bacilli, and Clostridia were scattered across in the right side. AMF, *Cyanobacterium*, and *Rickettsia* were located in the far-right side.

## Future Work

Microorganisms contain a number of heterogeneous alternative sigma factors that are selectively induced in response to environmental stress [38]. They not only provide functionally specialized RNA polymerase subpopulations, but are also involved in regulating the expression of a set of target genes, or regulon [39–40]. In contrast to sigma factors, regulatory systems governing heterogeneous alternative ribosome subpopulations in response to environmental stress remain largely unknown. Since multiple heterogeneous rRNA genes within a single isolate do not necessarily correlate with the extent of heterogeneity of functionally specialized ribosomes, sequence comparison of rRNA genes and ribosomal coding genes within a single isolate, as well as among closely related organisms, will be required in order to study their influence on adaptation of microorganism [41–42].

A codon is an attribute of a set of codes based on which transcriptional and translational regulators produce a gene product from a nucleotide sequence. Codon usage and codon context have been documented to play various important roles in these processes. If there are multiple types of the heterogeneous alternative regulators, there may be multiple sets of codes. Trimer usage biases of the Genus Specific DB were calculated based on the CDS database within a genus without considering alternative regulators and regulons. We may need to further anatomize evolutionary forces acting on multiple consecutive codons into greater detail, which may increase the accuracy of taxonomic classification. Moreover, comparative studies on alternative regulator subpopulations may provide useful insights into the development of genetic markers with which we can detect changes in microbial community structures in response to environmental stress (Figure S7). It may lead to new perspectives and strategies for improving the analysis of metagenome data, especially AMF inoculant field data sampled from highly stressful environments.

## Authors’ contributions

KJE designed the program and implemented it using the Java programming language. CA gave advice on developing scoring methods. HM provided knowledge on AMF experiments, the goals of the program, information on recent studies in AMF research, and helped to draft the manuscript. All authors read and approved the final manuscript.

## Competing interests

The authors have declared no competing interests.

## Acknowledgments

The authors gratefully acknowledge AFE (Éducation et de l’Enseignement supérieur Quebec), FESP (Faculté des études supérieures et postdoctorales de l’UdeM), and IRBV (Institut de Recherche en Biologie Végétale de l’Université de Montréal) for awarding scholarships to KJE. The authors gratefully acknowledge insightful comments from BachirIffis, David Walsh, Etienne Yergeau, Franck Stefani, Ivan de la Providencia, Jesse Shapiro, Sylvie Hamel, and Yves Terrat. We also thank Andrew Blakney for English editing.

## Supplementary material

**Figure S1 Histogram of the number of genera produced per answer** X-axis represents how many genera the trimer usage probability scoring method produced in an answer of a query sequence.

**Figure S2 Histogram of the rank of correct taxon group in answer** X-axis represents at which rank a correct taxon group occurred for the first time in an answer of a query sequence.

**Figure S3 Genera with similar trimer usage probability scores (Bacteria)** The number of occurrences of each genus included in the correct answers was counted in the bacterial genus CDS and non-CDS test sets. The figure shows the genera with relatively high occurrences. In general, the genera that had little difference in their trimer usage probability scores belonged to the same phylum.

**Figure S4 Genera with similar trimer usage probability scores (Fungi)** The number of occurrence of each genus included in the correct answers was counted in the fungal genus CDS and non-CDS test sets. The figure shows the genera with relatively high occurrences. *Clostridium* occurred the second highest in the AMF test set based on trimer usage probability scoring method. Both Agaricomycotina and Pezizomycotina belong to Dikarya, and the former occurred the second highest in the answers of the latter, and vice versa.

**Figure S5 Scatter plot of correct prediction proportion and log10 (inverse of P value score) A.** The mean of the correct prediction proportions was calculated per number of matching Three Codon DNA 9-mers in the bacterial, the fungal, and the AMF CDS test sets. For example, 22 sequences had 97 matching Three Codon DNA 9-mers and 20 of them were correctly classified in the bacterial test set; the mean was 0.91. **B.** and **C.** We calculated log10 (inverse of *P* value score) in order to show a relationship between the correct prediction proportion and *P* value score. For example, the multiplicative inverses of both *P* value scores, 1.0E-10 and 9.0E-10, were approximated to be 1.0E10, and base 10 logarithm of 1.0E10 was 10. B was based on the result from the trimer usage probability scoring method, while C was based on the result from the rank probability scoring method.

**Figure S6 Visualization of proximity relationships among 54 genera using MDS** The XY axis graph represents proximity relationships among 54 genera based on trimer usage biases of Three Codon DNA 9-mers identified in the presumably open reading frame of the example sequence. Genera belonging to the same taxonomic group in 13 taxon groups are indicated by the same background color.

**Figure S7 Heterogeneous regulator subpopulations within a single isolate** The symbols and question marks in the figure indicate the following questions from left to right. Under an optimal growth condition, do alternative regulators transcribe/translate genes? Does sigma factor regulate the expression of functionally specialized ribosomal rRNA and protein coding genes? Do heterogeneous regulator subpopulations produce structurally different gene products? Under environmental stress, do major regulators transcribe/translate genes?

**Table S1 Total number of the bacterial genes per genus**

**Table S2 Total number of the fungal genes and introns per genus**

**Table S3 Correct taxon group proportion in an answer in the bacterial test sets**

**Table S4 Correct taxon group proportion in an answer in the fungal test sets**

**Table S5 Frequency of the number of genera produced per answer**

**Table S6 Frequency of the rank of correct taxon group in answer**

**Table S7 Percentage of the other group in answers**

**Table S8 Correlation between the correct prediction proportion and the number of matching Three Codon DNA 9-mers**

**Table S9 Correlation between the correct prediction proportion of the trimer usage probability scoring method and *P* value score**

**Table S10 Correlation between the correct prediction proportion of the rank probability scoring method and *P* value score**

**Table S11 Relationship between the correct prediction proportion of the trimer usage probability scoring method and *P* value score in quartiles**

**Table S12 Relationship between the correct prediction proportion of the rank probability scoring method and *P* value score in quartiles**

## Notes

http://www.fungalsesame.org/

