## Supplementary Tables 1-12 and Figures S1-S7 for "SeSaMe: Metagenome Sequence Classification of Arbuscular Mycorrhizal Fungi Associated Microorganisms"

### Supplementary material

#### Kang et al. Genomics, Proteomiocs & Bioinformatics

**Supplementary Table 1  Total number of bacterial genes per genus**

| Bacterial genus | Total number of genes | Bacterial genus | Total number of genes |
|---|---|---|---|
| *Acidithiobacillus* | 12252 | *Microbacterium* | 3676 |
| *Acidobacterium* | 7924 | *Micrococcus* | 2236 |
| *Agrobacterium* | 22773 | *Myxococcus* | 22658 |
| *Anabaena* | 16055 | *Nitrobacter* | 7448 |
| *Azorhizobium* | 4717 | *Nitrosococcus* | 9744 |
| *Azotobacter* | 15105 | *Nitrosomonas* | 11481 |
| *Bacillus* | 392788 | *Nitrosospira* | 2805 |
| *Bdellovibrio* | 9939 | *Nocardia* | 19851 |
| *Beijerinckia* | 3784 | *Nostoc* | 16970 |
| *Bradyrhizobium* | 38418 | *Oscillatoria* | 12156 |
| *Caulobacter* | 17132 | *Pseudanabaena* | 3854 |
| *Clostridium* | 168122 | *Pseudomonas* | 321496 |
| *Cyanobacterium* | 6268 | *Pseudonocardia* | 6797 |
| *Desulfotomaculum* | 21547 | *Rhizobium* | 52249 |
| *Desulfovibrio* | 51438 | *Rhodobacter* | 20963 |
| *Erwinia* | 31931 | *Rickettsia* | 46400 |
| *Frankia* | 29711 | *Shewanella* | 102711 |
| *Geobacter* | 34729 | *Sinorhizobium* | 65703 |
| *Klebsiella* | 69640 | *Sphingomonas* | 15561 |
| *Kocuria* | 2356 | *Streptomyces* | 149826 |
| *Leuconostoc* | 16970 | *Variovorax* | 18984 |
| *Mesorhizobium* | 30388 | *Xanthomonas* | 65650 |
| *Methylococcus* | 2960 | | |

**Supplementary Table 2  Total number of fungal genes and introns per genus**

| Fungal genus | Total number of genes | Total number of introns |
|---|---|---|
| AMF | 21929 | 52385 |
| *Aspergillus* | 19695 | 38513 |
| *Cenococcum* | 14748 | 27036 |
| *Cryptococcus* | 13174 | 68068 |
| *Mycosphaerella* | 13107 | 33856 |
| *Oidiodendron* | 16703 | 32542 |
| *Phanerochaete* | 10048 | 48688 |
| *Scleroderma* | 21012 | 65184 |
| *Sebacina* | 15312 | 58256 |

**Supplementary Table 3  Correct taxon group proportion in an answer in bacterial test sets**

| Genus | CDS | | Non-CDS | | Genus | CDS | | Non-CDS | |
|---|---|---|---|---|---|---|---|---|---|
| | Mean | SD | Mean | SD | | Mean | SD | Mean | SD |
| *Acidithiobacillus* | 0.691 | 0.327 | 0.552 | 0.344 | *Microbacterium* | 0.948 | 0.166 | 0.892 | 0.229 |
| *Acidobacterium* | 0.825 | 0.303 | 0.616 | 0.379 | *Micrococcus* | 0.979 | 0.096 | 0.857 | 0.261 |
| *Agrobacterium* | 0.860 | 0.236 | 0.646 | 0.317 | *Myxococcus* | 0.912 | 0.216 | 0.562 | 0.317 |
| *Anabaena* | 0.762 | 0.311 | 0.745 | 0.299 | *Nitrobacter* | 0.911 | 0.187 | 0.767 | 0.295 |
| *Azorhizobium* | 0.922 | 0.188 | 0.708 | 0.294 | *Nitrosococcus* | 0.632 | 0.344 | 0.577 | 0.351 |
| *Azotobacter* | 0.788 | 0.287 | 0.792 | 0.298 | *Nitrosomonas* | 0.776 | 0.343 | 0.719 | 0.384 |
| *Bacillus* | 0.758 | 0.334 | 0.610 | 0.322 | *Nitrosospira* | 0.826 | 0.297 | 0.722 | 0.351 |
| *Bdellovibrio* | 0.886 | 0.247 | 0.715 | 0.358 | *Nocardia* | 0.850 | 0.254 | 0.755 | 0.285 |
| *Beijerinckia* | 0.834 | 0.279 | 0.662 | 0.341 | *Nostoc* | 0.927 | 0.169 | 0.770 | 0.305 |
| *Bradyrhizobium* | 0.928 | 0.175 | 0.696 | 0.289 | *Oscillatoria* | 0.830 | 0.297 | 0.696 | 0.357 |
| *Caulobacter* | 0.897 | 0.203 | 0.815 | 0.269 | *Pseudanabaena* | 0.939 | 0.184 | 0.643 | 0.344 |
| *Clostridium* | 0.864 | 0.252 | 0.792 | 0.285 | *Pseudomonas* | 0.840 | 0.250 | 0.664 | 0.327 |
| *Cyanobacterium* | 0.960 | 0.150 | 0.736 | 0.328 | *Pseudonocardia* | 0.972 | 0.127 | 0.911 | 0.202 |
| *Desulfotomaculum* | 0.782 | 0.308 | 0.662 | 0.304 | *Rhizobium* | 0.896 | 0.191 | 0.673 | 0.336 |
| *Desulfovibrio* | 0.592 | 0.330 | 0.504 | 0.317 | *Rhodobacter* | 0.934 | 0.190 | 0.687 | 0.267 |
| *Erwinia* | 0.841 | 0.279 | 0.585 | 0.339 | *Rickettsia* | 0.904 | 0.234 | 0.823 | 0.290 |
| *Frankia* | 0.907 | 0.201 | 0.841 | 0.277 | *Shewanella* | 0.773 | 0.336 | 0.715 | 0.337 |
| *Geobacter* | 0.770 | 0.299 | 0.594 | 0.344 | *Sinorhizobium* | 0.855 | 0.239 | 0.776 | 0.304 |
| *Klebsiella* | 0.893 | 0.236 | 0.746 | 0.331 | *Sphingomonas* | 0.917 | 0.181 | 0.757 | 0.267 |
| *Kocuria* | 0.967 | 0.130 | 0.885 | 0.238 | *Streptomyces* | 0.953 | 0.158 | 0.848 | 0.262 |
| *Leuconostoc* | 0.838 | 0.282 | 0.594 | 0.296 | *Variovorax* | 0.938 | 0.195 | 0.684 | 0.337 |
| *Mesorhizobium* | 0.876 | 0.214 | 0.729 | 0.304 | *Xanthomonas* | 0.904 | 0.215 | 0.751 | 0.319 |
| *Methylococcus* | 0.796 | 0.289 | 0.698 | 0.346 | **Mean** | **0.87** | **0.25** | **0.72** | **0.32** |

*Note*: After genera in an answer were converted to 13 taxon groups, the proportion of the correct taxon group was calculated per sequence in genus test set. The mean and the standard deviation of correct taxon group proportions are shown in the table.

#### Supplementary Table 4  Correct taxon group proportion in an answer in fungal test sets

| Genus | CDS | | Non-CDS | | Genus | CDS | | Non-CDS | |
|---|---|---|---|---|---|---|---|---|---|
| | Mean | SD | Mean | SD | | Mean | SD | Mean | SD |
| AMF | 0.706 | 0.302 | 0.689 | 0.317 | *Oidiodendron* | 0.385 | 0.280 | 0.326 | 0.233 |
| *Aspergillus* | 0.450 | 0.294 | 0.367 | 0.258 | *Phanerochaete* | 0.593 | 0.297 | 0.608 | 0.261 |
| *Cenococcum* | 0.467 | 0.276 | 0.429 | 0.282 | *Scleroderma* | 0.556 | 0.244 | 0.563 | 0.248 |
| *Cryptococcus* | 0.668 | 0.280 | 0.523 | 0.265 | *Sebacina* | 0.605 | 0.272 | 0.499 | 0.226 |
| *Mycosphaerella* | 0.650 | 0.309 | 0.457 | 0.255 | **Mean** | **0.567** | **0.298** | **0.498** | **0.28** |

*Note*: After genera in an answer were converted to 13 taxon groups, the proportion of the correct taxon group was calculated per sequence in genus test set. The mean and the standard deviation of correct taxon group proportions are shown in the table.

#### Supplementary Table 5  Frequency of the number of genera produced per answer

| | **1** | **2** | **3** | **4** | **5** | **6** | **7** | **8** | **9** | **10** | **11** | **12** | **Sum** |
|---|---|---|---|---|---|---|---|---|---|---|---|---|---|
| Bact. CDS: Correct | 2160 | 627 | 361 | 210 | 115 | 56 | 33 | 17 | 2 | 4 | 1 | 1 | 3587 |
| Bact. CDS: Incorrect | 281 | 189 | 143 | 126 | 80 | 49 | 23 | 16 | 6 | 0 | 0 | 0 | 913 |
| Fungi CDS: Correct | 170 | 125 | 121 | 87 | 90 | 50 | 23 | 10 | 2 | 0 | 0 | 0 | 678 |
| Fungi CDS: Incorrect | 87 | 62 | 41 | 14 | 9 | 6 | 1 | 1 | 0 | 1 | 0 | 0 | 222 |
| | | | | | | | | | | | | **Total** | 5400 |
| Bact. Non-CDS: Correct | 1358 | 539 | 418 | 242 | 163 | 108 | 67 | 37 | 18 | 14 | 5 | 1 | 2970 |
| Bact. Non-CDS: Incorrect | 521 | 363 | 222 | 165 | 144 | 58 | 38 | 15 | 3 | 0 | 1 | 0 | 1530 |
| Fungi non-CDS: Correct | 121 | 130 | 121 | 154 | 101 | 63 | 45 | 10 | 3 | 1 | 1 | 0 | 750 |
| Fungi non-CDS: Incorrect | 52 | 50 | 26 | 14 | 5 | 3 | 0 | 0 | 0 | 0 | 0 | 0 | 150 |
| | | | | | | | | | | | | **Total** | 5400 |

*Note*: The table shows frequencies of how many genera the trimer usage probability score method produced in an answer of a query sequence in correct and incorrect results in bacterial and fungal CDS and non-CDS test sets. Data for Supplementary Figure 1.

**Supplementary Table 6  Frequency of the rank of correct taxon group in answer**

| | 0 | 1 | 2 | 3 | 4 | 5 | 6 | 7 | 8 | 9 | 10 | Sum | Total |
|---|---|---|---|---|---|---|---|---|---|---|---|---|---|
| Bact. CDS | 3218 | 208 | 86 | 33 | 29 | 9 | 1 | 3 | 0 | 0 | 0 | 3587 | 4500 |
| Bact. Non-CDS | 2260 | 368 | 178 | 75 | 38 | 28 | 9 | 5 | 5 | 2 | 2 | 2970 | 4500 |
| Fungi CDS | 422 | 138 | 76 | 25 | 10 | 6 | 0 | 1 | 0 | 0 | 0 | 678 | 900 |
| Fungi non-CDS | 453 | 177 | 60 | 36 | 19 | 5 | 0 | 0 | 0 | 0 | 0 | 750 | 900 |

*Note*: The table shows frequency of at which rank the trimer usage probability score method produced a correct taxon group for the first time in an answer of a query sequence in correct results in bacterial and fungal CDS and non-CDS test sets. Data for Supplementary Figure 2.

**Supplementary Table 7  Percentage of the other group in answers**

| | Correct prediction percentages (genus) | The other group in answers | AMF in answers |
|---|---|---|---|
| Bact. CDS | 3185/4500: 71% | 197/3185: 6% | 119/4500: 3% |
| Bact. non-CDS | 2238/4500: 50% | 489/2238: 22% | 229/4500: 5% |
| Fung. CDS | 589/900: 65% | 93/589: 16% | 6/800: 1% |
| Fung. non-CDS | 655/900: 73% | 157/655: 24% | 10/800: 1% |
| AMF CDS | 49/100: 49% | 24/49: 49% | |
| AMF non-CDS | 72/100: 72% | 35/72: 49% | |

*Note*: The column- The other group in answers- indicates the percentage of fungal and bacterial group in answers of bacterial and fungal test sets, respectively, while it indicates the percentage of bacterial group in case of AMF test sets. The column- AMF in answers- indicates the percentage of AMF in answers of bacterial and fungal test sets.

**Supplementary Table 8 Correlation between correct prediction and the number of matching three codon DNA 9-mers**

| Bacteria | | Fungi | | AMF | |
|---|---|---|---|---|---|
| **No. of matched three codons** | **Correct prediction proportion** | **No. of matched three codons** | **Correct prediction proportion** | **No. of matched three codons** | **Correct prediction proportion** |
| 34 | 0.5 | 30 | 1 | 48 | 0.33 |
| 38 | 0.33 | 38 | 0.6 | 49 | 0.71 |
| 40 | 0.14 | 42 | 0.33 | 50 | 0.5 |
| 42 | 0.13 | 43 | 0.33 | 51 | 1 |
| 43 | 0.04 | 44 | 0.22 | 52 | 0.67 |
| 44 | 0.44 | 45 | 0.14 | 53 | 1 |
| 45 | 0.15 | 46 | 0.27 | 54 | 0.33 |
| 46 | 0.35 | 47 | 0.27 | 55 | 0.5 |
| 47 | 0.31 | 48 | 0.32 | 56 | 1 |
| 48 | 0.47 | 49 | 0.25 | 58 | 0.5 |
| 49 | 0.53 | 50 | 0.44 | 59 | 0.33 |
| 50 | 0.55 | 51 | 0.58 | 60 | 0.33 |
| 51 | 0.43 | 52 | 0.5 | 61 | 0.43 |
| 52 | 0.56 | 53 | 0.4 | 62 | 0.5 |
| 53 | 0.46 | 54 | 0.3 | 63 | 0.67 |
| 54 | 0.49 | 55 | 0.4 | 64 | 1 |
| 55 | 0.61 | 56 | 0.4 | 65 | 0.5 |
| 56 | 0.56 | 57 | 0.29 | 68 | 1 |
| 57 | 0.56 | 58 | 0.29 | 69 | 1 |
| 58 | 0.55 | 59 | 0.36 | 70 | 0.67 |
| 59 | 0.58 | 60 | 0.39 | 76 | 1 |
| 60 | 0.6 | 61 | 0.31 | 78 | 0.33 |
| 61 | 0.62 | 62 | 0.36 | 79 | 1 |
| 62 | 0.58 | 63 | 0.25 | 80 | 1 |
| 63 | 0.53 | 64 | 0.39 | 82 | 1 |
| 64 | 0.57 | 65 | 0.42 | 83 | 0.5 |
| 65 | 0.55 | 66 | 0.18 | 84 | 1 |
| 66 | 0.6 | 67 | 0.26 | 85 | 0.33 |
| 67 | 0.56 | 68 | 0.35 | 87 | 1 |
| 68 | 0.66 | 69 | 0.53 | 88 | 1 |
| 69 | 0.65 | 70 | 0.31 | 91 | 1 |
| 70 | 0.7 | 71 | 0.11 | 93 | 1 |
| 71 | 0.5 | 72 | 0.4 | | |
| 72 | 0.56 | 73 | 0.39 | | |
| 73 | 0.61 | 74 | 0.5 | | |
| 74 | 0.62 | 75 | 0.4 | | |
| 75 | 0.66 | 76 | 0.25 | | |
| 76 | 0.68 | 77 | 0.39 | | |

| | | | |
|---|---|---|---|
| 77 | 0.59 | 78 | 0.58 |
| 78 | 0.71 | 79 | 0.64 |
| 79 | 0.64 | 80 | 0.42 |
| 80 | 0.69 | 81 | 0.36 |
| 81 | 0.69 | 82 | 0.57 |
| 82 | 0.64 | 83 | 0.27 |
| 83 | 0.76 | 84 | 0.47 |
| 84 | 0.62 | 86 | 0.6 |
| 85 | 0.66 | 87 | 0.33 |
| 86 | 0.71 | 88 | 0.27 |
| 87 | 0.79 | 89 | 0.5 |
| 88 | 0.62 | 90 | 0.36 |
| 89 | 0.65 | 91 | 0.57 |
| 90 | 0.78 | 92 | 0.75 |
| 91 | 0.65 | 93 | 0.5 |
| 92 | 0.76 | 94 | 0.57 |
| 93 | 0.79 | 95 | 0.5 |
| 94 | 0.73 | 96 | 0.8 |
| 95 | 0.84 | 97 | 1 |
| 96 | 0.82 | | |
| 97 | 0.91 | | |

*Note*: Data for Supplementary Figure 5.A.

**Supplementary Table 9 Correlation between correct prediction of the trimer usage probability score method and *P* value score**

| **Bacteria** | | **Fungi** | | **AMF** | |
|---|---|---|---|---|---|
| **$Log_{10}$ (Inverse of *P* value score)** | **Correct prediction proportion** | **$Log_{10}$ (Inverse of *P* value score)** | **Correct prediction proportion** | **$Log_{10}$ (Inverse of *P* value score)** | **Correct prediction proportion** |
| 4 | 0 | 4 | 0 | 6 | 0 |
| 5 | 0.0833 | 5 | 0 | 8 | 0 |
| 6 | 0.0879 | 6 | 0.21 | 9 | 0 |
| 7 | 0.116 | 7 | 0.314 | 10 | 0.333 |
| 8 | 0.152 | 8 | 0.236 | 11 | 0.125 |
| 9 | 0.206 | 9 | 0.349 | 12 | 0 |
| 10 | 0.232 | 10 | 0.277 | 13 | 0.333 |
| 11 | 0.338 | 11 | 0.424 | 14 | 0.333 |
| 12 | 0.457 | 12 | 0.22 | 15 | 0.25 |
| 13 | 0.447 | 13 | 0.333 | 16 | 0.2 |
| 14 | 0.562 | 14 | 0.393 | 17 | 0 |
| 15 | 0.613 | 15 | 0.377 | 18 | 1 |
| 16 | 0.609 | 16 | 0.465 | 19 | 1 |
| 17 | 0.678 | 17 | 0.388 | 20 | 0.666 |

| | | | | | |
|---|---|---|---|---|---|
| 18 | 0.715 | 18 | 0.406 | 21 | 1 |
| 19 | 0.698 | 19 | 0.47 | 22 | 1 |
| 20 | 0.746 | 20 | 0.555 | 23 | 0.8 |
| 21 | 0.787 | 21 | 0.642 | 24 | 0.5 |
| 22 | 0.818 | 22 | 0.166 | 25 | 0.666 |
| 23 | 0.804 | 23 | 0.857 | 26 | 0.833 |
| 24 | 0.809 | 24 | 0.75 | 27 | 1 |
| 25 | 0.837 | 25 | 0.333 | 28 | 0.333 |
| 26 | 0.822 | 26 | 1 | 29 | 0.5 |
| 27 | 0.863 | 27 | 0.857 | 30 | 1 |
| 28 | 0.912 | 28 | 0.666 | 31 | 1 |
| 29 | 0.864 | 29 | 0.5 | 32 | 0.75 |
| 30 | 0.862 | 30 | 0.666 | 34 | 0 |
| 31 | 0.845 | 32 | 0.5 | 35 | 0.5 |
| 32 | 0.94 | 36 | 1 | 36 | 1 |
| 33 | 0.944 | 37 | 0 | 37 | 1 |
| 34 | 0.925 | | | 38 | 1 |
| 35 | 0.933 | | | 39 | 1 |
| 36 | 0.903 | | | 40 | 0 |
| 37 | 0.948 | | | 41 | 1 |
| 38 | 0.875 | | | 43 | 0 |
| 39 | 0.966 | | | 44 | 1 |
| 40 | 0.972 | | | 45 | 1 |
| 41 | 0.925 | | | 46 | 1 |
| 42 | 0.809 | | | 48 | 1 |
| 43 | 0.952 | | | 52 | 1 |
| 44 | 0.941 | | | 55 | 1 |
| 45 | 1 | | | | |
| 46 | 0.888 | | | | |
| 47 | 0.875 | | | | |
| 48 | 1 | | | | |
| 49 | 1 | | | | |
| 50 | 0.9 | | | | |
| 51 | 1 | | | | |
| 52 | 1 | | | | |
| 53 | 1 | | | | |
| 54 | 1 | | | | |
| 55 | 1 | | | | |
| 57 | 1 | | | | |
| 58 | 1 | | | | |
| 62 | 1 | | | | |
| 63 | 1 | | | | |

*Note*: The mean of the correct prediction proportions per $\log_{10}$ (Inverse of *P* value score) was calculated based on the first ranked genus with the highest probability score in the result from the trimer usage probability score method applied to bacterial, fungal, and AMF CDS test sets. Data for Supplementary Figure 5.B.

**Supplementary Table 10 Correlation between correct prediction of the rank probability score method and *P* value score**

| Bacteria | | Fungi | | AMF | |
|---|---|---|---|---|---|
| **$Log_{10}$ (Inverse of *P* value score)** | **Correct prediction proportion** | **$Log_{10}$ (Inverse of *P* value score)** | **Correct prediction proportion** | **$Log_{10}$ (Inverse of *P* value score)** | **Correct prediction proportion** |
| 5 | 0.166 | 5 | 0 | 6 | 0 |
| 6 | 0.325 | 6 | 0.315 | 7 | 0 |
| 7 | 0.386 | 7 | 0.363 | 8 | 0 |
| 8 | 0.445 | 8 | 0.565 | 9 | 0.25 |
| 9 | 0.612 | 9 | 0.74 | 10 | 0 |
| 10 | 0.676 | 10 | 0.762 | 11 | 0.142 |
| 11 | 0.717 | 11 | 0.674 | 12 | 0.166 |
| 12 | 0.768 | 12 | 0.767 | 13 | 0.285 |
| 13 | 0.821 | 13 | 0.805 | 14 | 0.6 |
| 14 | 0.828 | 14 | 0.847 | 15 | 0.3 |
| 15 | 0.881 | 15 | 0.871 | 16 | 0.5 |
| 16 | 0.929 | 16 | 0.763 | 17 | 0.666 |
| 17 | 0.922 | 17 | 0.794 | 18 | 1 |
| 18 | 0.939 | 18 | 0.933 | 19 | 0.75 |
| 19 | 0.952 | 19 | 0.833 | 20 | 0.666 |
| 20 | 0.916 | 20 | 0.8 | 21 | 0 |
| 21 | 0.964 | 21 | 0.857 | 22 | 1 |
| 22 | 0.965 | 22 | 1 | 23 | 0.333 |
| 23 | 0.961 | 23 | 1 | 24 | 1 |
| 24 | 0.989 | 24 | 1 | 25 | 1 |
| 25 | 0.95 | 25 | 0.5 | 26 | 0 |
| 26 | 0.942 | 26 | 0 | 29 | 1 |
| 27 | 0.98 | 29 | 1 | 30 | 1 |
| 28 | 0.96 | 30 | 1 | 31 | 0.5 |
| 29 | 0.975 | 41 | 1 | 32 | 1 |
| 30 | 0.941 | | | 34 | 1 |
| 31 | 0.941 | | | 35 | 1 |
| 32 | 0.923 | | | | |
| 33 | 1 | | | | |
| 34 | 1 | | | | |
| 35 | 1 | | | | |
| 36 | 1 | | | | |
| 37 | 1 | | | | |
| 38 | 1 | | | | |
| 39 | 1 | | | | |
| 40 | 1 | | | | |
| 42 | 1 | | | | |
| 43 | 1 | | | | |
| 47 | 1 | | | | |
| 52 | 1 | | | | |

*Note*: The mean of the correct prediction proportions per $log_{10}$ (Inverse of *P* value score) was calculated based on the first ranked genus with the highest probability score in the result from the rank probability score method applied to bacterial, fungal, and AMF CDS test sets. Data for Supplementary Figure 5.C.

**Supplementary Table 11 Relationship between correct prediction of the trimer usage probability score method and *P* value score in quartiles**

| Genus | 0 - 25th percentile | | | 26 - 50th percentile | | | 51 - 75th percentile | | | 76 - 100th percentile | | |
|---|---|---|---|---|---|---|---|---|---|---|---|---|
| | Range | Mean | SD | Range | Mean | SD | Range | Mean | SD | Range | Mean | SD |
| *Acidithiobacillus* | 5-8 | 0.151 | 0.18 | 9-13 | 0.41 | 0.217 | 14-19 | 0.461 | 0.402 | 20-32 | 0.6 | 0.547 |
| *Acidobacterium* | 6-10 | 0.276 | 0.18 | 11-15 | 0.564 | 0.223 | 16-20 | 0.62 | 0.073 | 21-28 | 0.875 | 0.306 |
| *Agrobacterium* | 4-9 | 0.045 | 0.0622 | 10-14 | 0.425 | 0.139 | 15-19 | 0.733 | 0.278 | 20-30 | 0.444 | 0.455 |
| *Anabaena* | 5-13 | 0.157 | 0.329 | 14-22 | 0.327 | 0.321 | 23-32 | 0.529 | 0.425 | 33-51 | 0.85 | 0.337 |
| *Azorhizobium* | 7-15 | 0.333 | 0.388 | 16-23 | 0.75 | 0.277 | 24-31 | 0.975 | 0.0707 | 32-43 | 1 | 0 |
| *Azotobacter* | 6-12 | 0.276 | 0.34 | 13-20 | 0.783 | 0.357 | 21-27 | 0.959 | 0.107 | 28-41 | 1 | 0 |
| *Bacillus* | 5-12 | 0.182 | 0.227 | 13-21 | 0.416 | 0.333 | 22-30 | 0.679 | 0.22 | 31-47 | 0.833 | 0.25 |
| *Bdellovibrio* | 6-11 | 0.205 | 0.186 | 12-18 | 0.669 | 0.308 | 19-25 | 0.821 | 0.144 | 26-37 | 0.928 | 0.188 |
| *Beijerinckia* | 5-9 | 0.08 | 0.109 | 10-15 | 0.473 | 0.279 | 16-21 | 0.906 | 0.0924 | 22-34 | 1 | 0 |
| *Bradyrhizobium* | 4-11 | 0.178 | 0.237 | 12-19 | 0.667 | 0.237 | 20-26 | 0.857 | 0.196 | 27-40 | 0.937 | 0.176 |
| *Caulobacter* | 6-13 | 0.328 | 0.37 | 14-22 | 0.781 | 0.213 | 23-31 | 0.982 | 0.0505 | 32-42 | 1 | 0 |
| *Clostridium* | 6-20 | 0.424 | 0.389 | 21-32 | 0.83 | 0.211 | 33-45 | 1 | 0 | 47-63 | 0.972 | 0.0962 |
| *Cyanobacterium* | 7-19 | 0.233 | 0.344 | 20-30 | 0.903 | 0.205 | 31-41 | 0.893 | 0.238 | 42-54 | 1 | 0 |
| *Desulfotomaculum* | 5-10 | 0.201 | 0.178 | 11-16 | 0.42 | 0.291 | 17-22 | 0.48 | 0.43 | 23-38 | 0.69 | 0.365 |
| *Desulfovibrio* | 6-10 | 0.253 | 0.31 | 11-15 | 0.276 | 0.145 | 16-20 | 0.23 | 0.338 | 21-30 | 0.0833 | 0.204 |
| *Erwinia* | 5-10 | 0.163 | 0.146 | 11-16 | 0.488 | 0.289 | 17-22 | 0.877 | 0.142 | 23-34 | 1 | 0 |
| *Frankia* | 5-12 | 0.272 | 0.309 | 13-20 | 0.72 | 0.197 | 21-27 | 0.45 | 0.326 | 28-46 | 0.375 | 0.443 |
| *Geobacter* | 6-9 | 0.243 | 0.204 | 10-14 | 0.464 | 0.232 | 15-18 | 0.69 | 0.359 | 19-28 | 0.85 | 0.223 |
| *Klebsiella* | 5-10 | 0.236 | 0.409 | 11-17 | 0.642 | 0.135 | 18-23 | 0.979 | 0.051 | 24-32 | 1 | 0 |
| *Kocuria* | 6-15 | 0.275 | 0.415 | 17-27 | 0.75 | 0.403 | 29-38 | 0.987 | 0.0395 | 39-54 | 0.977 | 0.0753 |
| *Leuconostoc* | 7-14 | 0.166 | 0.288 | 15-23 | 0.854 | 0.242 | 24-31 | 0.979 | 0.0589 | 32-44 | 1 | 0 |
| *Mesorhizobium* | 5-10 | 0.116 | 0.139 | 11-16 | 0.546 | 0.345 | 17-22 | 0.682 | 0.205 | 23-31 | 0.597 | 0.395 |
| *Methylococcus* | 6-10 | 0.14 | 0.219 | 11-15 | 0.711 | 0.309 | 16-20 | 0.763 | 0.152 | 21-30 | 0.934 | 0.106 |
| *Microbacterium* | 7-16 | 0.0937 | 0.265 | 17-26 | 0.922 | 0.171 | 27-34 | 0.933 | 0.128 | 37-50 | 1 | 0 |
| *Micrococcus* | 10-19 | 0.15 | 0.253 | 20-30 | 0.91 | 0.156 | 31-40 | 0.983 | 0.0527 | 41-55 | 1 | 0 |
| *Myxococcus* | 6-16 | 0.461 | 0.373 | 17-25 | 0.869 | 0.182 | 26-34 | 1 | 0 | 35-49 | 1 | 0 |
| *Nitrobacter* | 6-10 | 0.155 | 0.175 | 11-15 | 0.467 | 0.193 | 16-20 | 0.5 | 0.204 | 21-40 | 0.611 | 0.443 |
| *Nitrosococcus* | 4-8 | 0.0333 | 0.0745 | 9-14 | 0.12 | 0.138 | 15-19 | 0.762 | 0.146 | 20-27 | 0.777 | 0.403 |
| *Nitrosomonas* | 5-10 | 0.203 | 0.211 | 11-16 | 0.449 | 0.192 | 17-22 | 0.875 | 0.209 | 23-48 | 0.833 | 0.408 |
| *Nitrosospira* | 6-9 | 0.106 | 0.093 | 10-14 | 0.644 | 0.328 | 15-19 | 0.883 | 0.111 | 20-25 | 0.8 | 0.447 |
| *Nocardia* | 6-12 | 0.226 | 0.229 | 13-19 | 0.717 | 0.271 | 20-26 | 0.826 | 0.149 | 27-36 | 0.802 | 0.35 |
| *Nostoc* | 5-12 | 0.168 | 0.252 | 13-20 | 0.509 | 0.151 | 21-28 | 0.676 | 0.232 | 29-41 | 0.388 | 0.485 |
| *Oscillatoria* | 6-11 | 0.194 | 0.155 | 12-18 | 0.63 | 0.143 | 19-25 | 0.821 | 0.256 | 26-34 | 1 | 0 |
| *Pseudanabaena* | 6-12 | 0.161 | 0.203 | 13-19 | 0.778 | 0.246 | 20-26 | 0.952 | 0.125 | 27-41 | 0.875 | 0.353 |
| *Pseudomonas* | 7-12 | 0.255 | 0.389 | 13-18 | 0.691 | 0.196 | 19-24 | 0.885 | 0.18 | 25-35 | 0.773 | 0.368 |
| *Pseudonocardia* | 9-19 | 0.323 | 0.404 | 20-30 | 0.815 | 0.229 | 31-40 | 0.966 | 0.105 | 41-55 | 1 | 0 |
| *Rhizobium* | 5-9 | 0.125 | 0.19 | 10-15 | 0.222 | 0.178 | 16-21 | 0.376 | 0.287 | 22-28 | 0.805 | 0.305 |
| *Rhodobacter* | 7-14 | 0.216 | 0.357 | 15-22 | 0.848 | 0.217 | 23-30 | 0.921 | 0.175 | 32-48 | 1 | 0 |
| *Rickettsia* | 5-21 | 0.399 | 0.459 | 22-33 | 0.845 | 0.2 | 34-44 | 0.85 | 0.312 | 45-62 | 1 | 0 |
| *Shewanella* | 5-9 | 0.333 | 0.471 | 10-15 | 0.619 | 0.344 | 16-22 | 0.646 | 0.186 | 23-30 | 0.777 | 0.403 |
| *Sinorhizobium* | 4-10 | 0.16 | 0.158 | 11-15 | 0.288 | 0.208 | 16-20 | 0.616 | 0.273 | 21-30 | 0.7 | 0.447 |
| *Sphingomonas* | 6-13 | 0.135 | 0.274 | 14-21 | 0.689 | 0.327 | 22-29 | 0.907 | 0.202 | 30-40 | 1 | 0 |
| *Streptomyces* | 5-15 | 0.101 | 0.154 | 16-24 | 0.765 | 0.193 | 25-33 | 0.856 | 0.194 | 34-50 | 0.796 | 0.328 |
| *Variovorax* | 6-15 | 0.222 | 0.44 | 16-25 | 0.93 | 0.113 | 26-35 | 1 | 0 | 36-49 | 1 | 0 |
| *Xanthomonas* | 7-13 | 0.454 | 0.252 | 14-20 | 0.83 | 0.187 | 21-28 | 0.94 | 0.104 | 29-42 | 1 | 0 |
| AMF | 6-16 | 0.157 | 0.15 | 17-26 | 0.746 | 0.315 | 27-37 | 0.708 | 0.358 | 38-55 | 0.818 | 0.404 |

| | | | | | | | | | | | | |
|---|---|---|---|---|---|---|---|---|---|---|---|---|
| *Aspergillus* | 6-10 | 0.405 | 0.234 | 11-15 | 0.24 | 0.205 | 16-20 | 0.116 | 0.162 | 21-37 | 0.2 | 0.447 |
| *Cenococcum* | 6-10 | 0.165 | 0.205 | 11-15 | 0.331 | 0.158 | 16-20 | 0.133 | 0.217 | 21-32 | 0.25 | 0.418 |
| *Cryptococcus* | 6-10 | 0.228 | 0.435 | 11-16 | 0.292 | 0.285 | 17-23 | 0.475 | 0.404 | 24-32 | 0.833 | 0.408 |
| *Mycosphaerella* | 4-8 | 0.266 | 0.326 | 9-13 | 0.56 | 0.308 | 14-18 | 0.518 | 0.153 | 19-26 | 0.542 | 0.366 |
| *Oidiodendron* | 5-9 | 0.15 | 0.223 | 10-14 | 0.325 | 0.129 | 15-19 | 0.526 | 0.345 | 20-25 | 0.533 | 0.505 |
| *Phanerochaete* | 6-9 | 0.471 | 0.11 | 10-14 | 0.424 | 0.157 | 15-19 | 0.373 | 0.127 | 21-30 | 0.6 | 0.547 |
| *Scleroderma* | 5-9 | 0.04 | 0.0894 | 10-14 | 0.287 | 0.149 | 15-19 | 0.388 | 0.146 | 20-36 | 0.9 | 0.223 |
| *Sebacina* | 5-9 | 0.0833 | 0.117 | 10-14 | 0.357 | 0.153 | 15-19 | 0.558 | 0.275 | 20-28 | 1 | 0 |

*Note*: Range represents minimum and maximum of $\log_{10}$ (Inverse of *P* value score) values per quartile. After result data from each genus test set were divided into quartiles, the range of $\log_{10}$ (inverse of *P* value score) and the mean and the standard deviation of correct prediction proportions were calculated per quartile. The result data were based on trimer usage probability score method and the same as those for Supplementary Table 9 and Supplementary Figure 5.B.

**Supplementary Table 12  Relationship between correct prediction of the rank probability score method and *P* value score in quartiles**

| Genus | 0-25th percentiles | | | 26-50th percentiles | | | 51-75th percentiles | | | 76-100th percentiles | | |
|---|---|---|---|---|---|---|---|---|---|---|---|---|
| | Range | Mean | SD | Range | Mean | SD | Range | Mean | SD | Range | Mean | SD |
| *Acidithiobacillus* | 6-9 | 0.278 | 0.242 | 10-13 | 0.748 | 0.0505 | 14-17 | 0.968 | 0.0625 | 18-40 | 1 | 0 |
| *Acidobacterium* | 6-10 | 0.277 | 0.277 | 11-15 | 0.82 | 0.106 | 16-20 | 1 | 0 | 21-26 | 1 | 0 |
| *Agrobacterium* | 6-9 | 0.718 | 0.359 | 10-14 | 0.923 | 0.07 | 15-19 | 1 | 0 | 20-27 | 1 | 0 |
| *Anabaena* | 7-12 | 0.189 | 0.244 | 13-18 | 0.721 | 0.37 | 19-24 | 0.901 | 0.113 | 25-31 | 1 | 0 |
| *Azorhizobium* | 7-12 | 0.762 | 0.237 | 13-19 | 1 | 0 | 20-25 | 1 | 0 | 26-34 | 1 | 0 |
| *Azotobacter* | 6-11 | 0.567 | 0.343 | 12-17 | 0.894 | 0.117 | 18-24 | 1 | 0 | 25-43 | 0.833 | 0.408 |
| *Bacillus* | 5-11 | 0.244 | 0.308 | 12-18 | 0.77 | 0.226 | 19-24 | 0.777 | 0.194 | 25-37 | 1 | 0 |
| *Bdellovibrio* | 7-11 | 0.712 | 0.188 | 12-16 | 0.923 | 0.104 | 17-21 | 0.96 | 0.0894 | 22-27 | 1 | 0 |
| *Beijerinckia* | 5-10 | 0.6 | 0.383 | 11-15 | 0.9 | 0.173 | 16-20 | 1 | 0 | 21-34 | 1 | 0 |
| *Bradyrhizobium* | 7-11 | 0.712 | 0.209 | 12-16 | 0.95 | 0.0684 | 17-21 | 1 | 0 | 22-29 | 1 | 0 |
| *Caulobacter* | 7-11 | 0.689 | 0.317 | 12-16 | 0.971 | 0.0638 | 17-21 | 0.92 | 0.178 | 22-29 | 1 | 0 |
| *Clostridium* | 7-13 | 0.122 | 0.19 | 14-20 | 0.439 | 0.255 | 21-26 | 0.891 | 0.174 | 27-40 | 0.857 | 0.377 |
| *Cyanobacterium* | 11-17 | 0.516 | 0.273 | 18-24 | 0.823 | 0.262 | 25-31 | 0.976 | 0.0629 | 32-43 | 1 | 0 |
| *Desulfotomaculum* | 6-10 | 0.133 | 0.217 | 11-15 | 0.617 | 0.0981 | 16-20 | 0.893 | 0.153 | 22-35 | 1 | 0 |
| *Desulfovibrio* | 6-9 | 0.254 | 0.176 | 10-13 | 0.647 | 0.109 | 14-17 | 0.85 | 0.191 | 18-24 | 1 | 0 |
| *Erwinia* | 5-10 | 0.513 | 0.484 | 11-16 | 0.826 | 0.0847 | 17-22 | 1 | 0 | 23-30 | 1 | 0 |
| *Frankia* | 5-8 | 0.107 | 0.214 | 9-13 | 0.729 | 0.219 | 14-18 | 1 | 0 | 19-23 | 0.95 | 0.111 |
| *Geobacter* | 6-9 | 0.502 | 0.413 | 10-13 | 0.697 | 0.0762 | 14-17 | 1 | 0 | 18-22 | 0.9 | 0.223 |
| *Klebsiella* | 5-10 | 0.548 | 0.325 | 11-15 | 0.913 | 0.123 | 16-20 | 0.93 | 0.109 | 21-26 | 1 | 0 |
| *Kocuria* | 7-12 | 0.516 | 0.449 | 13-18 | 0.944 | 0.136 | 19-24 | 0.875 | 0.209 | 25-34 | 1 | 0 |
| *Leuconostoc* | 8-12 | 0.243 | 0.265 | 13-17 | 0.682 | 0.171 | 18-22 | 0.971 | 0.0638 | 23-27 | 1 | 0 |
| *Mesorhizobium* | 6-10 | 0.86 | 0.167 | 11-15 | 0.937 | 0.0908 | 16-20 | 1 | 0 | 21-28 | 1 | 0 |
| *Methylococcus* | 7-10 | 0.5 | 0.408 | 11-15 | 0.853 | 0.123 | 16-19 | 1 | 0 | 20-24 | 1 | 0 |
| *Microbacterium* | 7-11 | 0.4 | 0.418 | 12-17 | 0.877 | 0.113 | 18-23 | 1 | 0 | 24-32 | 1 | 0 |
| *Micrococcus* | 8-13 | 0.549 | 0.389 | 14-19 | 0.933 | 0.0831 | 20-24 | 1 | 0 | 25-35 | 1 | 0 |
| *Myxococcus* | 6-11 | 0.222 | 0.186 | 12-17 | 0.733 | 0.188 | 18-23 | 0.979 | 0.051 | 24-34 | 1 | 0 |
| *Nitrobacter* | 6-10 | 0.671 | 0.393 | 11-15 | 1 | 0 | 16-20 | 1 | 0 | 21-40 | 1 | 0 |
| *Nitrosococcus* | 5-8 | 0.583 | 0.3 | 9-13 | 0.752 | 0.11 | 14-17 | 0.937 | 0.125 | 18-24 | 1 | 0 |
| *Nitrosomonas* | 6-11 | 0.546 | 0.328 | 12-17 | 0.88 | 0.117 | 18-22 | 0.933 | 0.149 | 23-52 | 1 | 0 |
| *Nitrosospira* | 5-9 | 0.253 | 0.347 | 10-14 | 0.956 | 0.0603 | 15-19 | 1 | 0 | 20-26 | 1 | 0 |

| | | | | | | | | | | | | |
|---|---|---|---|---|---|---|---|---|---|---|---|---|
| *Nocardia* | 6-10 | 0.283 | 0.298 | 11-16 | 0.767 | 0.183 | 17-21 | 0.98 | 0.0447 | 22-35 | 1 | 0 |
| *Nostoc* | 8-12 | 0.511 | 0.5 | 13-17 | 0.662 | 0.197 | 18-22 | 0.9 | 0.136 | 23-31 | 0.916 | 0.204 |
| *Oscillatoria* | 6-10 | 0.336 | 0.232 | 11-16 | 0.886 | 0.137 | 17-21 | 1 | 0 | 22-29 | 1 | 0 |
| *Pseudanabaena* | 6-12 | 0.236 | 0.29 | 13-19 | 0.927 | 0.0922 | 20-26 | 0.984 | 0.0419 | 27-38 | 1 | 0 |
| *Pseudomonas* | 6-9 | 0.602 | 0.225 | 10-14 | 0.807 | 0.155 | 15-19 | 0.981 | 0.0406 | 20-28 | 1 | 0 |
| *Pseudonocardia* | 7-13 | 0.553 | 0.375 | 14-19 | 1 | 0 | 20-24 | 1 | 0 | 25-32 | 1 | 0 |
| *Rhizobium* | 6-9 | 0.493 | 0.359 | 10-14 | 0.918 | 0.0784 | 15-18 | 0.958 | 0.0833 | 19-25 | 1 | 0 |
| *Rhodobacter* | 6-11 | 0.726 | 0.405 | 12-18 | 0.947 | 0.0899 | 19-24 | 1 | 0 | 25-38 | 1 | 0 |
| *Rickettsia* | 8-17 | 0.199 | 0.247 | 18-25 | 0.67 | 0.242 | 26-33 | 0.651 | 0.342 | 34-47 | 1 | 0 |
| *Shewanella* | 5-9 | 0.25 | 0.204 | 10-14 | 0.708 | 0.136 | 15-18 | 0.947 | 0.0611 | 19-27 | 1 | 0 |
| *Sinorhizobium* | 5-8 | 0.875 | 0.25 | 9-13 | 0.923 | 0.104 | 14-18 | 0.98 | 0.0447 | 19-24 | 1 | 0 |
| *Sphingomonas* | 6-11 | 0.56 | 0.347 | 12-17 | 0.883 | 0.204 | 18-24 | 0.972 | 0.068 | 25-33 | 1 | 0 |
| *Streptomyces* | 5-10 | 0.139 | 0.219 | 11-16 | 0.863 | 0.141 | 17-22 | 1 | 0 | 23-32 | 1 | 0 |
| *Variovorax* | 7-12 | 0.291 | 0.367 | 13-18 | 0.715 | 0.172 | 19-24 | 0.986 | 0.034 | 25-31 | 1 | 0 |
| *Xanthomonas* | 7-11 | 0.516 | 0.207 | 12-17 | 0.839 | 0.202 | 18-23 | 1 | 0 | 25-39 | 1 | 0 |
| AMF | 6-11 | 0.0654 | 0.106 | 12-18 | 0.502 | 0.284 | 19-25 | 0.678 | 0.386 | 26-35 | 0.785 | 0.393 |
| *Aspergillus* | 5-8 | 0.398 | 0.377 | 9-12 | 0.87 | 0.0933 | 13-16 | 0.873 | 0.148 | 17-26 | 0.75 | 0.5 |
| *Cenococcum* | 6-9 | 0.625 | 0.25 | 10-13 | 0.82 | 0.208 | 14-17 | 0.703 | 0.216 | 18-41 | 0.72 | 0.414 |
| *Cryptococcus* | 6-9 | 0.339 | 0.282 | 10-14 | 0.61 | 0.286 | 15-18 | 0.825 | 0.236 | 19-25 | 0.8 | 0.447 |
| *Mycosphaerella* | 6-9 | 0.816 | 0.137 | 10-13 | 0.83 | 0.05 | 14-17 | 0.895 | 0.125 | 18-25 | 1 | 0 |
| *Oidiodendron* | 6-8 | 0.555 | 0.509 | 9-12 | 0.794 | 0.151 | 13-16 | 0.958 | 0.0833 | 17-22 | 0.816 | 0.213 |
| *Phanerochaete* | 5-7 | 0.154 | 0.135 | 8-11 | 0.488 | 0.213 | 12-15 | 0.78 | 0.182 | 16-21 | 0.775 | 0.262 |
| *Scleroderma* | 6-10 | 0.474 | 0.245 | 11-15 | 0.679 | 0.164 | 16-20 | 0.971 | 0.0638 | 21-30 | 1 | 0 |
| *Sebacina* | 7-9 | 0.466 | 0.416 | 10-13 | 0.69 | 0.183 | 14-17 | 0.843 | 0.119 | 18-23 | 1 | 0 |

*Note*: Range represents minimum and maximum of $\log_{10}$ (Inverse of $P$ value score) values per quartile. After result data from each genus test set were divided into quartiles, the range of $\log_{10}$ (inverse of $P$ value score) and the mean and the standard deviation of proportions of correct taxon group were calculated per quartile. The result data were based on rank probability score method and the same as those for Supplementary Table 10 and Supplementary Figure 5.C.

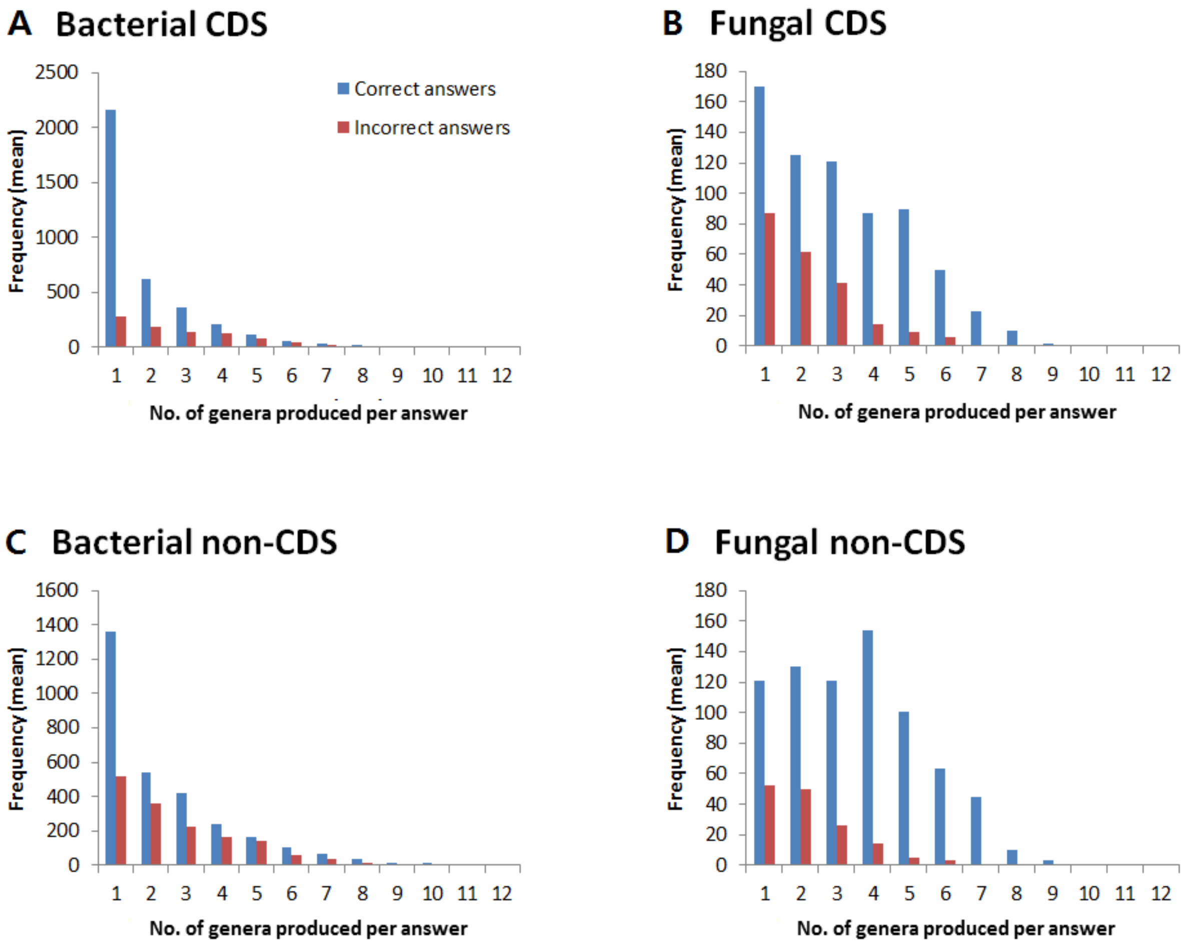


**Figure S1 Histogram of the number of genera produced per answer**

X-axis represents how many genera the trimer usage probability scoring method produced in an answer of a query sequence.

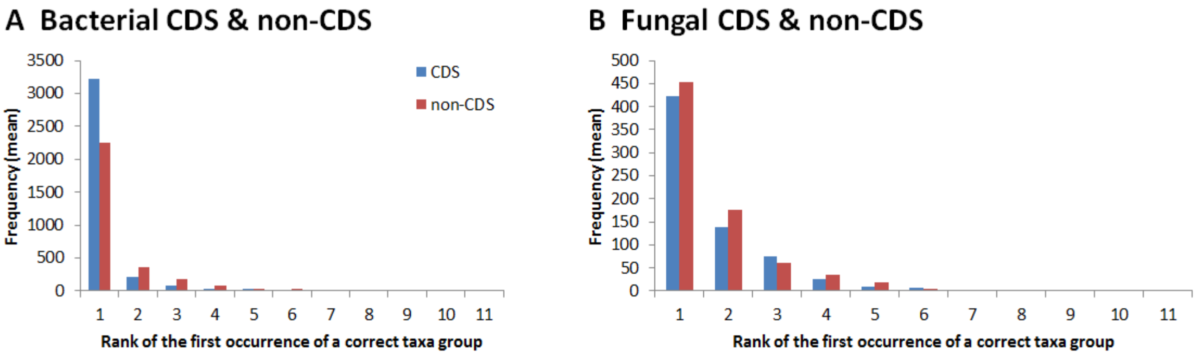


**Figure S2 Histogram of the rank of correct taxon group in answer**

X-axis represents at which rank a correct taxon group occurred for the first time in an answer of a query sequence.

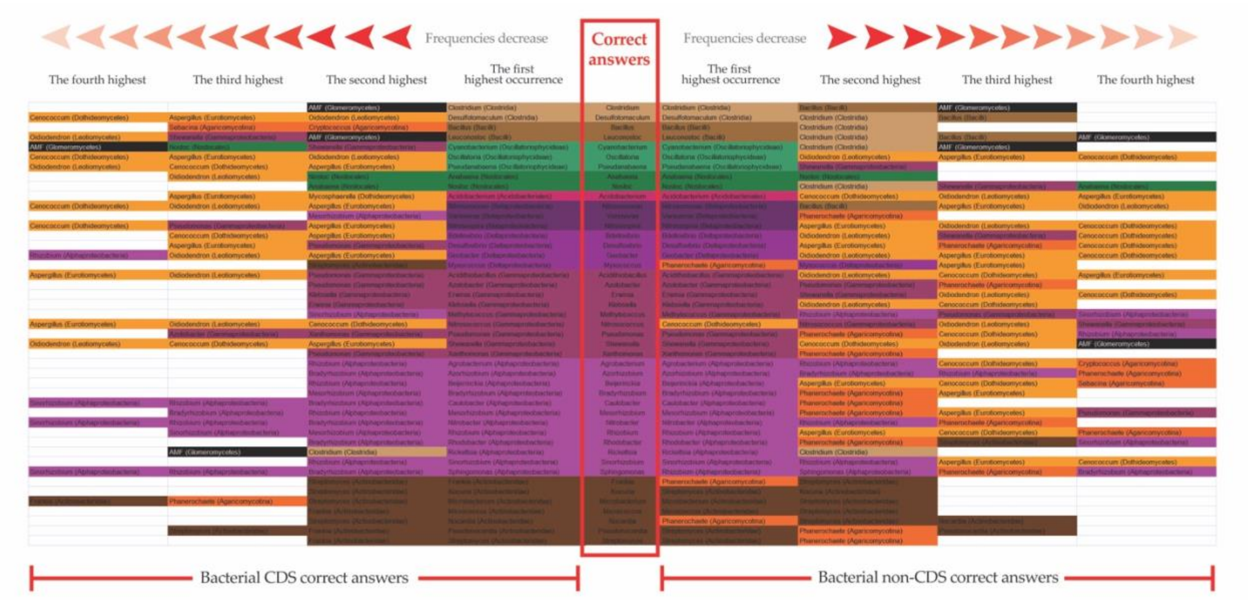


**Figure S3 Genera with similar trimer usage probability scores (Bacteria)**

The number of occurrences of each genus included in the correct answers was counted in the bacterial genus CDS and non-CDS test sets. The figure shows the genera with relatively high occurrences. In general, the genera that had little difference in their trimer usage probability scores belonged to the same phylum.

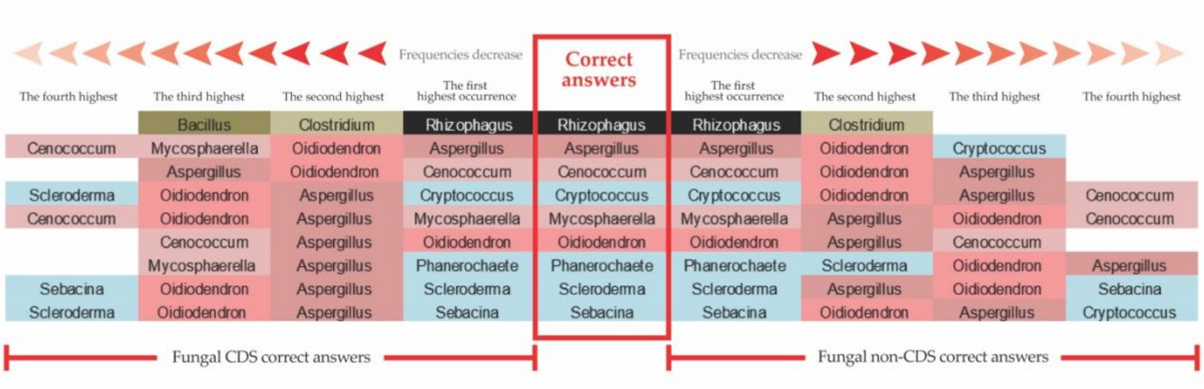


**Figure S4  Genera with similar trimer usage probability scores (Fungi)**

The number of occurrence of each genus included in the correct answers was counted in the fungal genus CDS and non-CDS test sets. The figure shows the genera with relatively high occurrences. *Clostridium* occurred the second highest in the AMF test set based on trimer usage probability scoring method. Both Agaricomycotina and Pezizomycotina belong to Dikarya, and the former occurred the second highest in the answers of the latter, and vice versa.

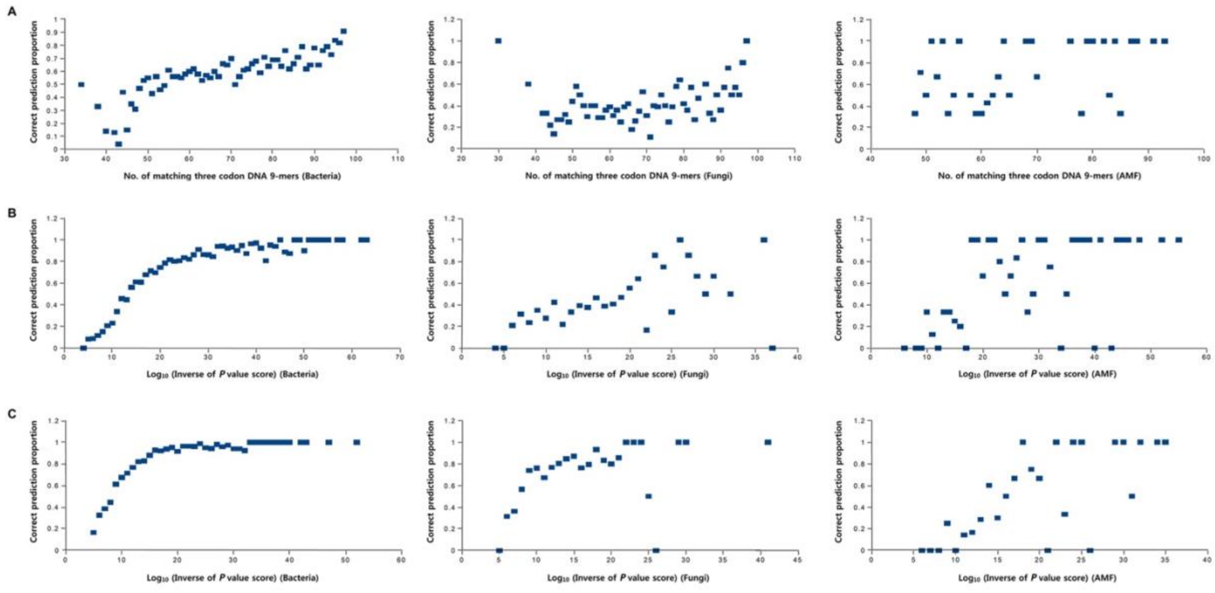


**Figure S5  Scatter plot of correct prediction proportion and log10 (inverse of *P* value score)**

**A.** The mean of the correct prediction proportions was calculated per number of matching Three Codon DNA 9-mers in the bacterial, the fungal, and the AMF CDS test sets. For example, 22 sequences had 97 matching Three Codon DNA 9-mers and 20 of them were correctly classified in the bacterial test set; the mean was 0.91. **B.** and **C.** We calculated $\log_{10}$ (inverse of *P* value score) in order to show a relationship between the correct prediction proportion and *P* value score. For example, the multiplicative inverses of both *P* value scores, 1.0E-10 and 9.0E-10, were approximated to be 1.0E10, and base 10 logarithm of 1.0E10 was 10. B was based on the result from the trimer usage probability scoring method, while C was based on the result from the rank probability scoring method.

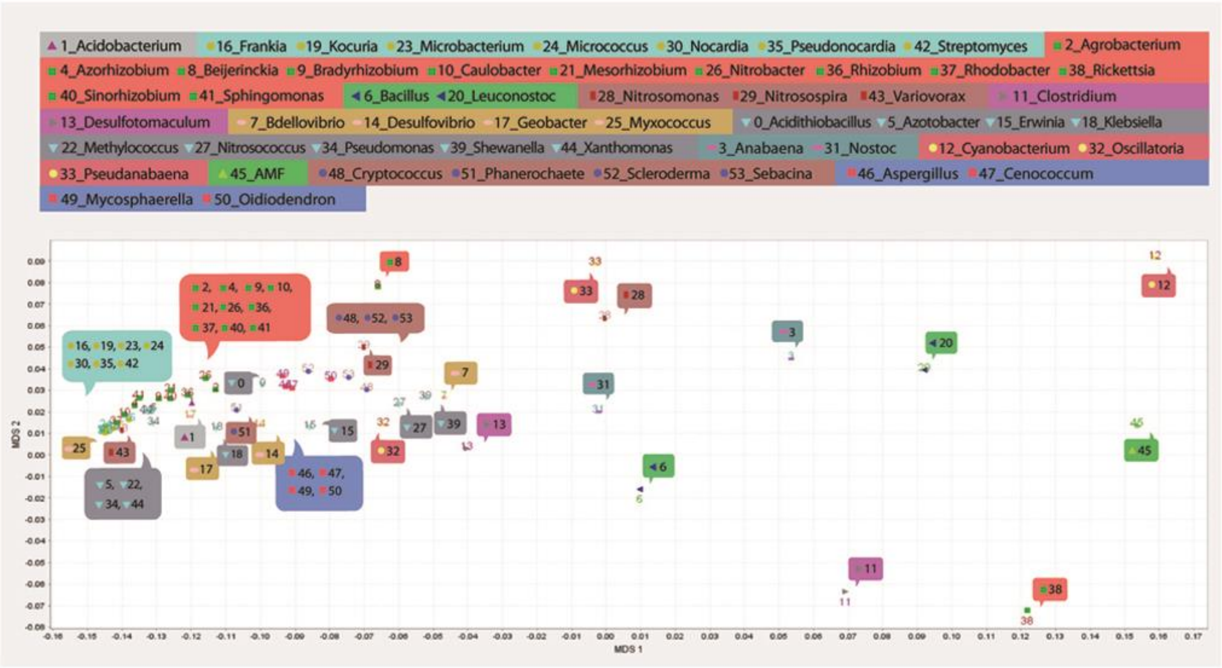


**Figure S6 Visualization of proximity relationships among 54 genera using MDS**

The XY axis graph represents proximity relationships among 54 genera based on trimer usage biases of Three Codon DNA 9-mers identified in the presumably open reading frame of the example sequence. Genera belonging to the same taxonomic group in 13 taxon groups are indicated by the same background color.

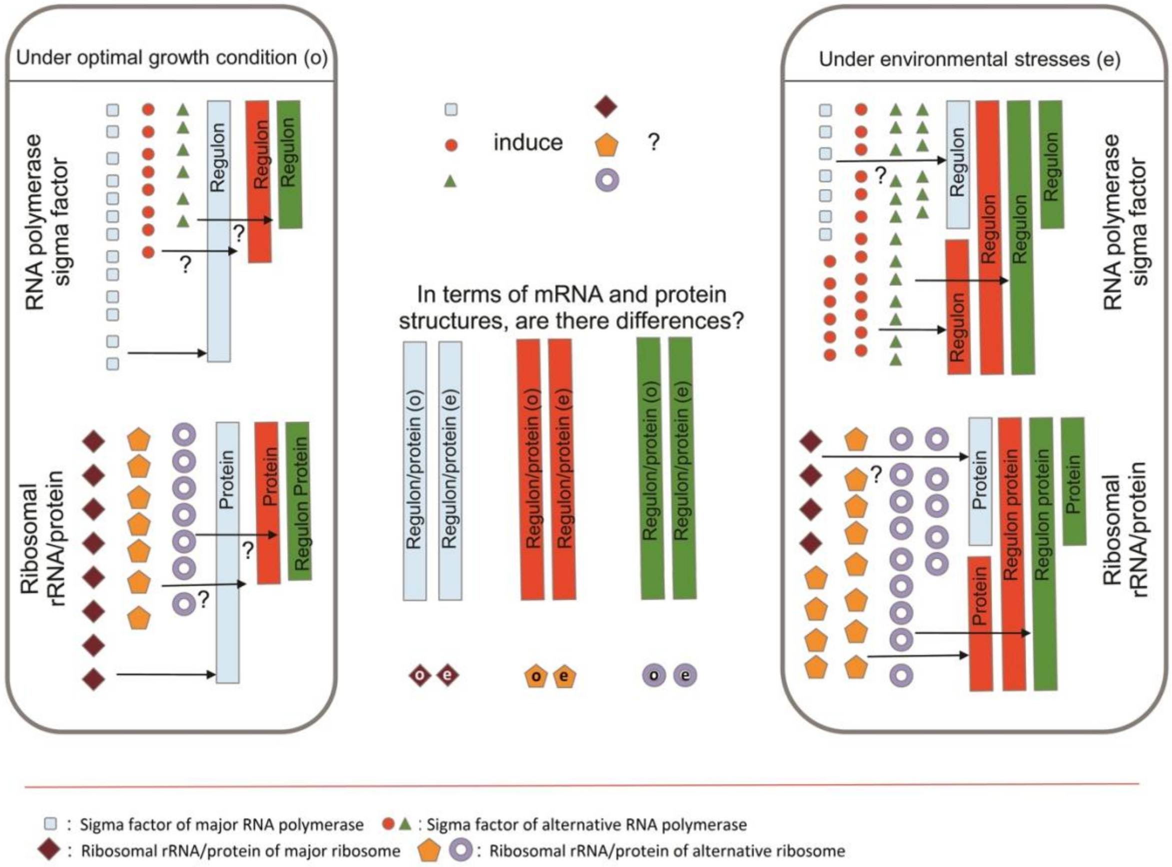


**Figure S7 Heterogeneous regulator subpopulations within a single isolate**

The symbols and question marks in the figure indicate the following questions from left to right. Under an optimal growth condition, do alternative regulators transcribe/translate genes? Does sigma factor regulate the expression of functionally specialized ribosomal rRNA and protein coding genes? Do heterogeneous regulator subpopulations produce structurally different gene products? Under environmental stress, do major regulators transcribe/translate genes?
